# The length of the expressed 3’ UTR is an intermediate molecular phenotype linking genetic variants to complex diseases

**DOI:** 10.1101/540088

**Authors:** Elisa Mariella, Federico Marotta, Elena Grassi, Stefano Gilotto, Paolo Provero

## Abstract

In the last decades, genome wide association studies (GWAS) have uncovered tens of thousands of associations between common genetic variants and complex diseases. However, these statistical associations can rarely be interpreted functionally and mechanistically. As the majority of the disease-associated variants are located far from coding sequences, even the relevant gene is often unclear. A way to gain insight into the relevant mechanisms is to study the genetic determinants of intermediate molecular phenotypes, such as gene expression and transcript structure. We propose a computational strategy to discover genetic variants affecting the relative expression of alternative 3’ untranslated region (UTR) isoforms, generated through alternative polyadenylation, a widespread post-transcriptional regulatory mechanism known to have relevant functional consequences. When applied to a large dataset in which whole genome and RNA sequencing data are available for 373 European individuals, 2,530 genes with alternative polyadenylation quantitative trait loci (apaQTL) were identified. We analyze and discuss possible mechanisms of action of these variants, and we show that they are significantly enriched in GWAS hits, in particular those concerning immune-related and neurological disorders. Our results point to an important role for genetically determined alternative polyadenylation in affecting predisposition to complex diseases, and suggest new ways to extract functional information from GWAS data.

## 1 Introduction

Understanding the relationship between human genotypes and phenotypes is one of the central goals of biomedical research. The first sequencing of the human genome [1, 2] and the following large-scale investigations of genetic differences between individuals by efforts such as the 1000 Genome Project Consortium [3] provided the foundation for the study of human genetics at the genome-wide level. Enrichment of trait-specific GWAS hits among apaQTL

Genome wide association studies (GWAS) examine common genetic variants to identify associations with complex traits, including common diseases. Long lists of genetic associations with disparate traits have been obtained, but their functional interpretation is far from being straightforward [4]. Indeed, because of linkage disequilibrium, GWAS identify genomic regions carrying multiple variants among which it is not possible to identify the causal ones without additional information. Furthermore, most loci identified in human GWAS are in non-coding regions, presumably exerting regulatory effects, but usually we do not known the identity of the affected gene or the molecular mechanism involved.

A possible way to gain insight into the mechanisms behind GWAS associations is to investigate the effect of genetic variants on intermediate molecular phenotypes, such as gene expression [5, 6]. Expression quantitative trait loci (eQTL) are genomic regions carrying one or more genetic variants affecting gene expression. Besides their intrinsic interest in understanding the control of gene expression, eQTL studies can be exploited for the interpretation of GWAS results, helping to prioritize likely causal variants and supporting the formulation of mechanistic hypotheses about the links between genetic variants and diseases.

Recent studies have shown that genetic variants acting on the whole RNA processing cascade are at least equally common as, and largely independent from, those that affect transcriptional activity, and that they can be a major driver of phenotypic variability in humans [7]. Therefore it is important to identify the genetic variants associated to transcript structure, including splicing and alternative UTR isoforms, besides those affecting transcriptional levels, and different approaches have been proposed to this end [8, 9]. From these studies it emerges in particular that genetic variants frequently determine changes in the length of the expressed 3’ UTRs. In addition, genome-wide analyses specifically focused on alternative splicing have been performed [10, 11].

Polyadenylation is one of the post-transcriptional modifications affecting pre-mRNAs in the nucleus and involves two steps: the cleavage of the transcript and the addition of a poly(A) tail [12, 13]. The most important regulatory elements involved are the polyadenylation signal (PAS) and other cis-elements, usually located within the 3’UTR, but multiple and diversified regulatory mechanisms have been described [14, 15]. The PAS is recognized by the cleavage and polyadenylation specificity factor (CPSF) that, together with other protein complexes, induces the cleavage of the transcript in correspondence of the downstream poly(A) site. The large majority of human genes have multiple poly(A) sites, so that alternative polyadenylation (APA) is a widespread phenomenon contributing to the diversification of the human transcriptome through the generation of alternative mature transcripts with different 3’ ends. Such transcripts are translated into identical proteins, but protein level, localization, and even interactions can depend on the 3’ end of the transcript [16].

APA events have been grouped into classes based on the location of the alternative poly(A) site and the type of change determined by their differential usage [12]. In this work we have taken into consideration only the simplest and most frequent mode (tandem 3’UTR APA), in which two poly(A) sites located within the same terminal exon, one in a proximal and one in a distal position, produce transcripts that differ only in the length of the 3’UTR. Such variation in 3’UTR length can have an important functional impact, for example by affecting the binding of microRNAs and RNA binding proteins and thus transcript abundance, translation and localization. Moreover, APA regulation is strongly tissue- and cell type-dependent [17, 18, 19, 20, 21] and several examples are known of altered APA regulation associated to human diseases [7, 22].

How genetic variants influence APA has not been comprehensively investigated in a large human population yet. A recent analysis of whole genome sequencing (WGS) data from [8] found hundreds of common single nucleotide polymorphisms (SNPs) causing the alteration or degradation of motifs that are similar to the canonical PAS [23], but did not extend the analysis to other possible mechanisms. Other studies found strong associations between genetic variants and APA regulation [24, 25, 26, 8, 9, 27], but a systematic investigation based on a large number of samples and variants, specifically targeted to APA rather than generically to transcript structure, and unbiased in the choice of variants to examine, is not yet available.

Here we propose a new computational strategy for the genome-wide investigation of the influence of genetic variants on the expression of alternative 3’UTR isoforms in a large population. In particular, we analyzed WGS data paired with standard RNA-Seq data obtained in 373 European (EUR) individuals [8]. Statistically, our approach is analogous to methods commonly implemented in eQTL mapping analysis and it aims to overcome the limitations illustrated above for the specific purpose of correlating variants to 3’ UTR isoforms.

A central task, preliminary to the analysis of genetic variants, is thus the quantification of the alternative 3’UTR isoforms. Various strategies have been implemented to this end, from custom analysis pipelines for microarray data [28], to the development of next-generation sequencing technologies specifically targeted to the 3’ end of transcripts, such as the serial analysis of gene expression (SAGE) [20] and sequencing of APA sites (SAPAs) [21], allowing also the identification of previously unannotated APA sites.

More recently, tools able to capture APA events from standard RNA-Seq data have been developed. In general, these approaches can be divided into two categories: those that exploit previous annotation of poly(A) sites [29, 30], such the ones provided by PolyA_DB2 [31] and APASdb [32], and those that instead try to infer their location from the data [17]. Although the latter approach potentially allows analyzing also previously unannotated sites, the former leads to higher sensitivity [29, 30], and was thus preferred in this study. Undoubtedly, approaches based on standard RNA-Seq are not as powerful and accurate as technologies that specifically sequence the 3’ ends. However, they allow studying this phenomenon in an uncomparably larger number of samples and conditions, including the recently generated large-scale transcriptomic datasets of normal individuals that we use in this work.

## 2 Results

### 2.1 Genetic variants affect the relative expression of alternative 3’UTR isoforms of thousands of genes

In order to investigate the effect of human genetic variants on the expression of alternative 3’UTR isoforms, we developed a computational approach similar to the one commonly used for eQTL analysis (Fig. 1). It was applied to a large dataset in which WGS data paired with RNA-Seq data are available for 373 European (EUR) individuals (GEUVADIS dataset [8]). A collection of known alternative poly(A) sites [31] was used, together with a compendium of human transcripts, to obtain an annotation of alternative 3’UTR isoforms that was then combined with RNA-Seq data in order to compute, for each gene, the expression ratio between short and long isoform (*m/M* value) in each individual.

**Figure 1:**
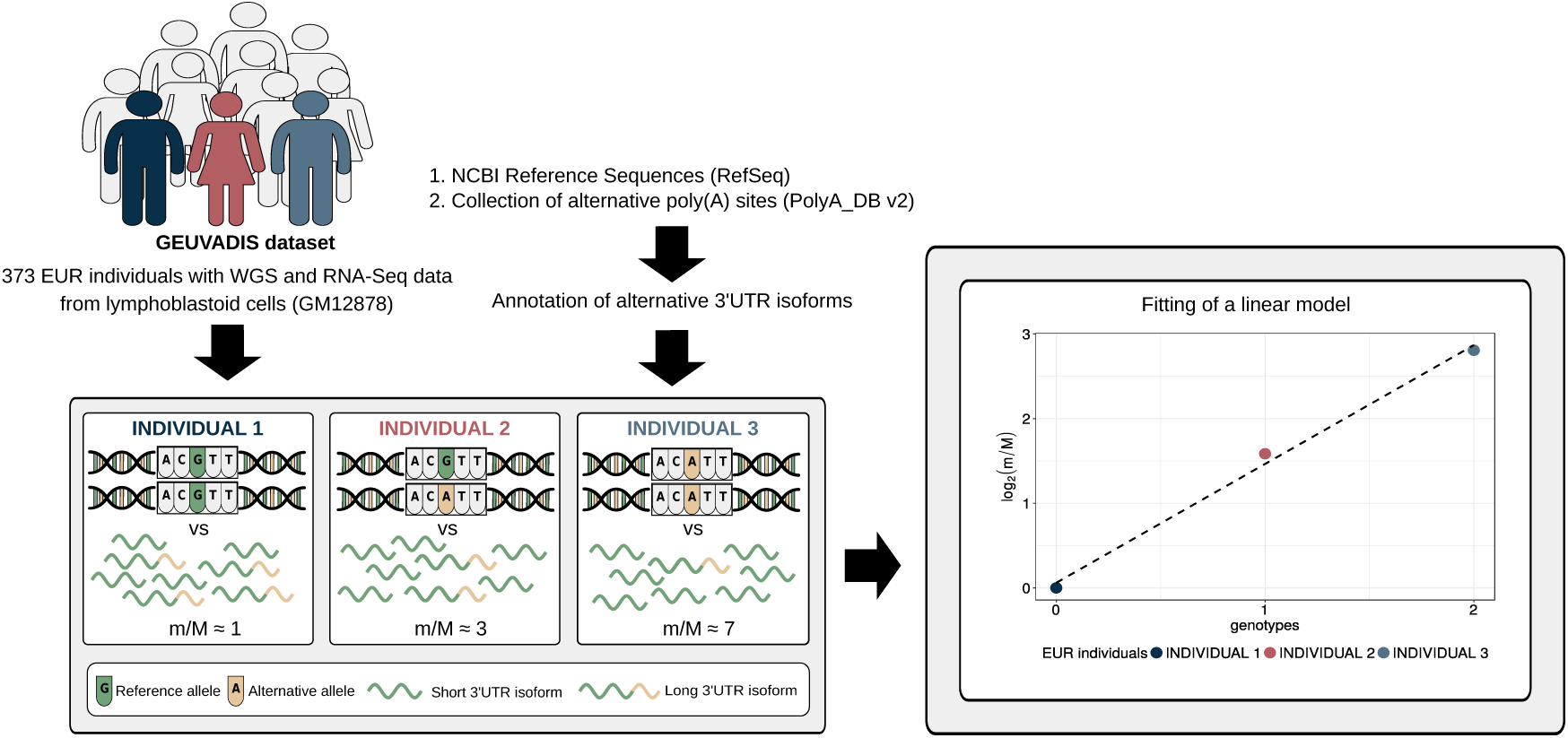
Schematic representation of the method. Genotypic data paired with RNA-Seq data from a large cohort of individuals are required to perform apaQTL mapping analysis. RNA-Seq data are exploited, together with an annotation of alternative 3’UTR isoforms, to compute for each gene the m/M value that is proportional to the ratio between the expression of its short and long 3’UTR isoforms. Then, the association between the m/M values of a gene and each nearby genetic variant is evaluated by linear regression. Genotypes are defined in the standard way: 0 means homozygous for the reference allele, 1 means heterozygous and 2 indicates the presence of two copies of the alternative allele.

Linear regression was then used to identify associations between the *m/M* values of each gene and the genetic variants within a cis-window including the gene itself and all sequence located within 1Mbp from the transcription start site (TSS) or the transcription end site (TES). This led to the fitting of *∼*30 million linear models, involving *∼*6,300 genes and *∼*5.3 million variants. About 190,000 models, involving 2,530 genes and *∼*160,000 variants, revealed a significant association (Fig. 2, Tab. 1, Supplementary Files 3 and 4).

**Figure 2:**
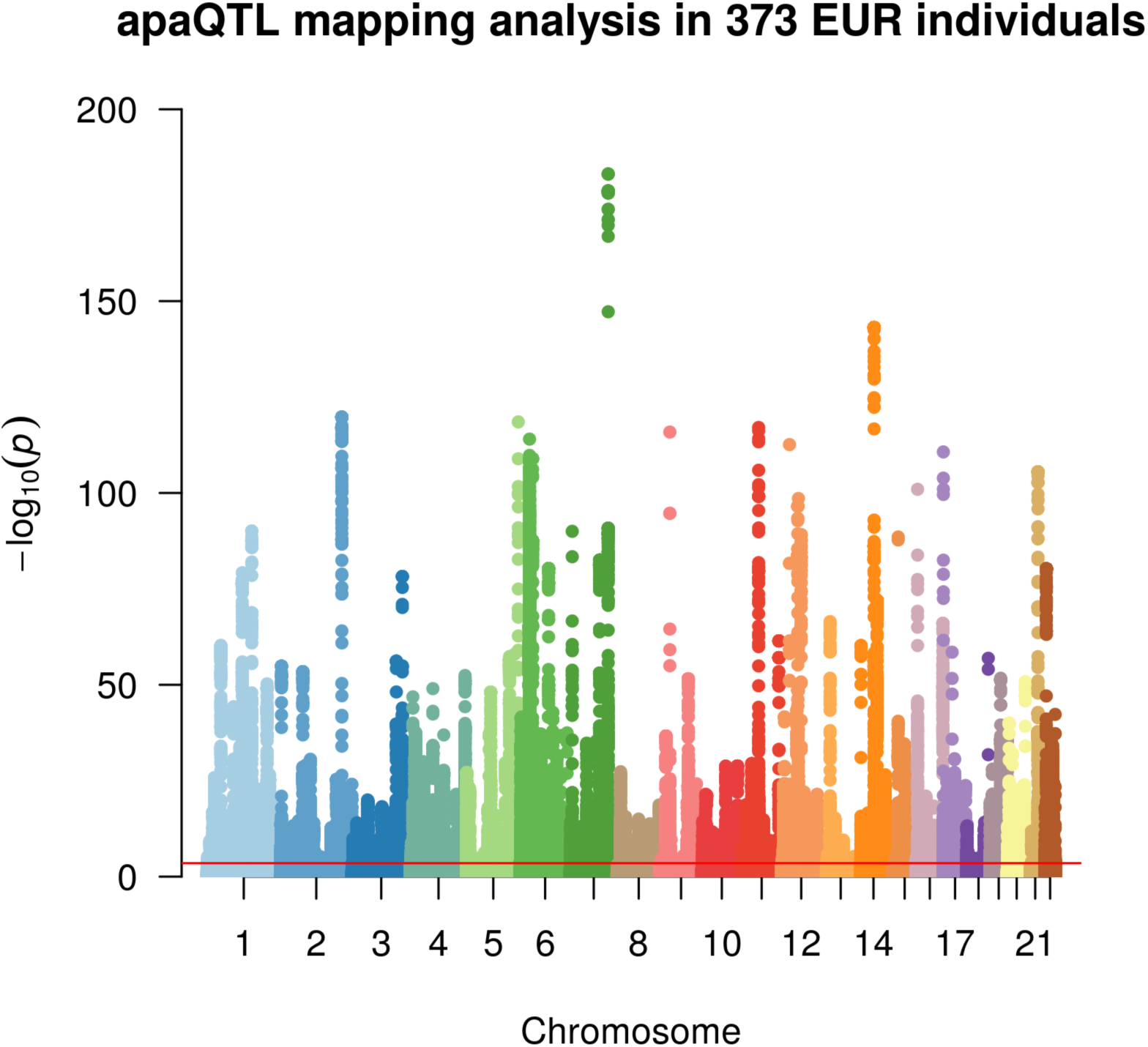
Manhattan plot illustrating the results of the apaQTL mapping analysis at the genome-wide level. For each fitted model, the-log_10_ P-value is shown according to the position of the tested genetic variant. The red line indicates the threshold for genome-wide statistical significance, after multiple-testing correction (nominal P-value < 3.1 *×* 10^*−*4^).

**Table 1:** Number of models, genes and variants.

|  | Total | Significant* |
| --- | --- | --- |
| Models | 30,136,480 | 192,715 |
| Genes | 6,256 | 2,530 |
| Variants | 5,309,860 | 160,223 |

\* corrected empirical P-value $< 0.05$

Our set of significant genes shows only moderate overlap with genes for which eQTLs or transcript ratio QTLs (trQTLs) were reported in Ref. [8] from the same data (Fig. 3). Alternative polyadenylation can result in changes in gene expression levels as a consequence of the isoform-dependent availability of regulatory elements affecting the stability of transcripts, such as microRNA binding sites [13]. In this case, apaQTLs should also be eQTLs. However, APA may also have effects that do not imply changes in expression levels, including the modulation of mRNA translation rates [33, 34] and localization [35], and protein cytoplasmic localization [36]. Similarly, a complete overlap with trQTLs is not expected, because they were identified by taking into account all the annotated alternative transcripts of a gene including alternative splicing and transcription initiation: The identification of apaQTLs for several genes for which trQTLs were not identified suggests that focusing on a specific class of transcript structure allows higher sensitivity.

**Figure 3:**
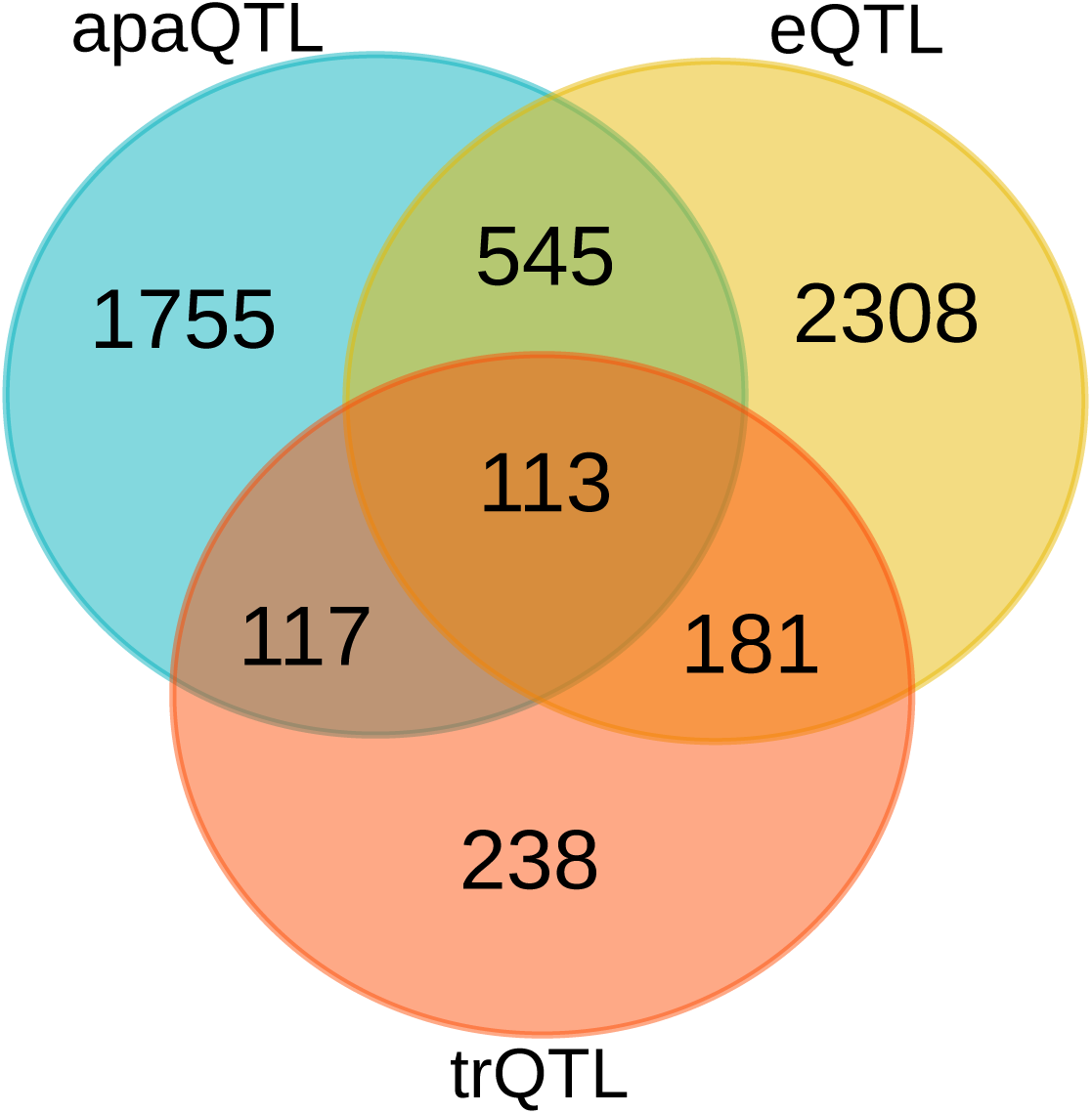
Overlap between genes with significant alternative polyadenylation QTL (apaQTL), expression QTL (eQTL) and transcript ratio QTL (trQTL).

These results show that a large number of genetic determinants of alternative polyadenylation can be inferred from the analysis of standard RNA-Seq data paired with the genotypic characterization on the same individuals.

### 2.2 apaQTLs are preferentially located within active genomic regions

Just like eQTLs, we expect apaQTLs be located within genomic regions that are active in the relevant cell type (lymphoblastoid cells for our data). In order to verify this hypothesis, we superimposed the apaQTLs to the ChromHMM annotation of the human genome for the GM12878 cell line [37], and used logistic regression, as detailed in the Methods, to determine the enrichment or depletion of apaQTLs for each chromatin state, expressed as an odds ratio (OR). As expected, significant ORs *>* 1 were obtained for active genomic regions, such as transcribed regions, promoters and enhancers, suggesting that genetic variants have a higher probability of being apaQTLs when they are located in active regions. Conversely, apaQTLs were depleted in repressed and inactive chromatin states. Similar results were obtained using broad chromatin states (Fig. 4), defined following [37], or all 15 chromatin states reported by ChromHMM (Supplementary Fig. 3).

**Figure 4:**
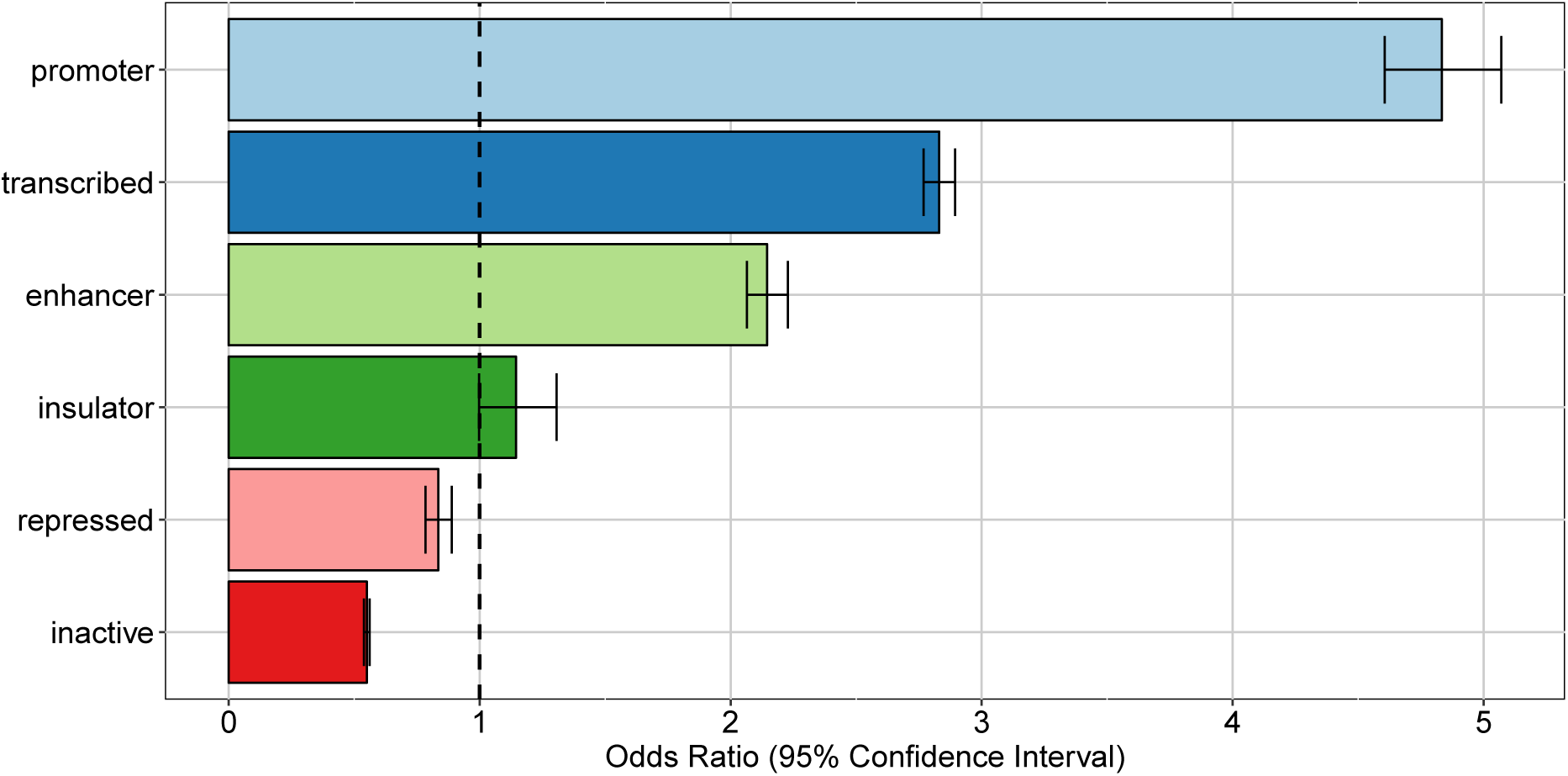
Enrichment of apaQTLs within broad chromatin states that were defined starting from the ChromHMM annotation. For each state, the OR obtained by logistic regression and its 95% CI are shown.

As a control, the same enrichment analysis was performed with the ChromHMM annotation obtained in a different cell type, namely normal human epithelial keratinocytes (NHEK). All NHEK active chromatin states showed a reduced enrichment in apaQTLs compared with GM1278, and regions repressed in NHEK cells actually showed significant enrichment of lymphoblastoid apaQTLs (Supplementary Fig. 4 and Supplementary Fig. 5). Taken together, these results show that genetic variants affecting alternative polyadenylation tend to be located in cell-type specific active chromatin regions.

In the following, we will divide apaQTLs in two classes: Intragenic apaQTLs are those located inside one of the genes whose isoform ratio we are able to analyze, while all other apaQTLs will be referred to as extragenic (note that these might be located inside a gene for which we are unable to perform the analysis, for one of the reasons explained in the Methods).

### 2.3 Intragenic apaQTLs are enriched in coding exons and 3’UTRs

Having established that genetic variants have a widespread influence of the expression of alternative 3’UTR isoforms, we turned to their putative mechanisms of action. First of all, we considered the distribution of intragenic apaQTLs among regions contributing to the mRNA vs. introns. As shown in Fig. 5, intragenic apaQTLs are enriched in coding exons and 3’ UTRs, and depleted in introns and 5’ UTRs. The depletion of introns suggests that most intragenic apaQTLs exert their regulatory role at the transcript level, e.g. by modulating the binding of trans-acting factors to the mRNA.

**Figure 5:**
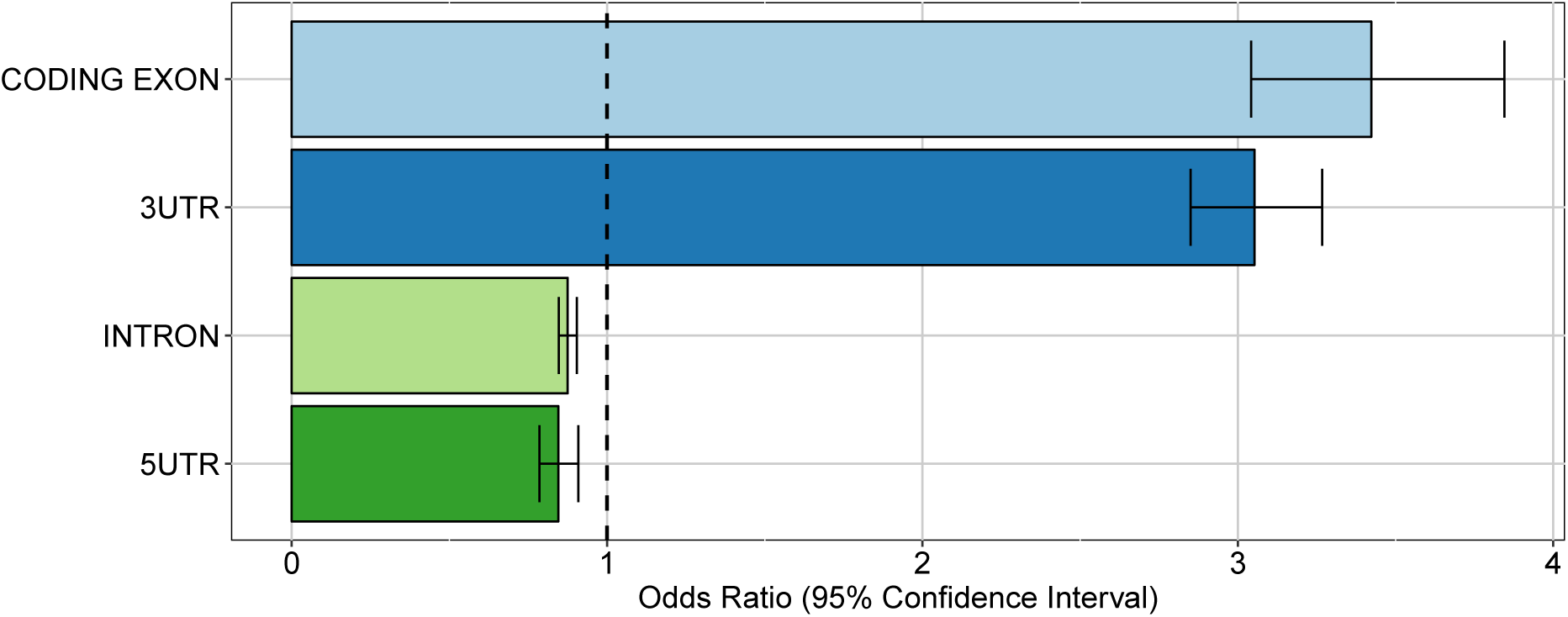
Enrichment of intragenic apaQTLs within coding and non-coding transcript regions. For each gene region, the OR obtained by logistic regression its 95% CI are shown.

Among mRNA regions, the enrichment of 3’ UTRs is expected, since these regions contain several elements involved in the regulation of both alternative polyadenylation and mRNA stability. The enrichment of coding exons could be ascribed to regulatory elements residing in these portions of the mRNAs, or to residual effects of linkage disequilibrium (LD) with variants located in the 3’ UTR, notwithstanding the LD pruning procedure implemented in the enrichment analysis (see Methods). Note that while several poly(A) sites are located upstream of the last exon [38], within both intronic sequences and internal exons, such sites were not taken into account in our analysis. Finally, the depletion of 5’ UTRs might be due to the distance of these elements from the polyadenylation loci, and to the fact that these regions are mostly involved in other regulatory mechanisms, such as translational regulation [39]. In the following, we examine in more detail three possible mechanisms by which intragenic apaQTLs could exert their action.

#### 2.3.1 Creation and destruction of PAS motifs

The first possibility is direct interference with the APA regulation, favoring the production of one of the two isoforms in individuals with a particular genotype. A comprehensive atlas of high-confidence PAS has been recently reported [40]. In addition to the canonical PAS motifs (AAUAAA and AUUAAA) it contains 10 previously known signals and 6 new motifs. Exploiting this resource, we were able to identify SNPs that cause the creation or the destruction of putative functional PAS motifs and, as expected, we found that they were enriched among apaQTLs (OR = 1.72, 95% confidence interval (CI) = 1.08 - 2.75, P-value = 0.0215). In total, 42 PAS-altering variants were found to be apaQTLs of the gene in which they reside. While expected, this result can be considered to validate our strategy.

A few examples are worth discussing in detail. SNP rs10954213 was shown by several studies [41, 24, 42] to determine the preferential production of the short isoform of the IRF5 transcription factor through the conversion of an alternative PAS motif (AAUGAA) into the canonical one (AAUAAA) in a proximal position within the 3’UTR. Consistently, we found that this variant is associated with higher prevalence of the short isoform (Fig. 6A,B). Moreover, the same variant was associated to higher risk of systemic lupus erythematosus (SLE), and higher IRF5 expression, that could be due to the loss of AU-rich elements (ARE) in the short transcript isoform [24]. Globally, these findings are in agreement with the known involvement of IRF5 in several pathways that are critical for the onset of SLE (Type I IFN production, M1 macrophage polarization, autoantibody production, and induction of apoptosis [43]).

**Figure 6:**
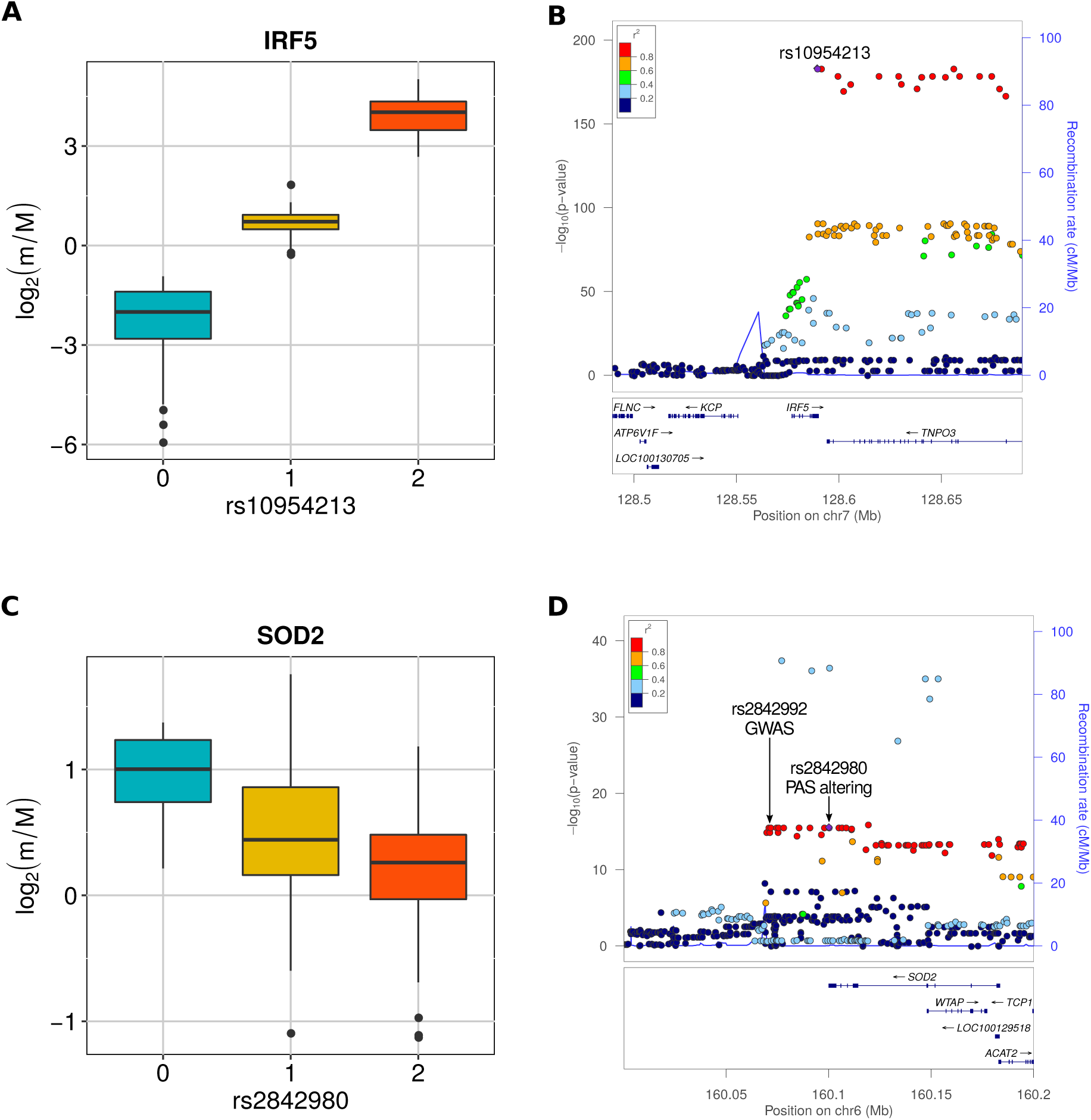
(A) Boxplot showing the variation of the log2-transformed m/M values obtained for IRF5 as a function of the genotype of the individuals for rs10954213. (B) Illustration of the results obtained for IRF5 in the genomic region around rs10954213 (100kb both upstream and downstream its genomic location). In the top panel each tested genetic variant was reported as a function of both its genomic coordinate and its association level with IRF5 (log10-transformed nominal P-value); the points color reflects the LD level (*R*^2^) between rs10954213 and each of the other genetic variants in the locus. The bottom panel shows the genes and their orientation in the locus. The figure was generated by LocusZoom [107]. (C) Boxplot showing the variation of the log2-transformed m/M values obtained for SOD2 as a function of the genotype of the individuals for rs2842980. (D) LocusZoom plot illustrating the results obtained for SOD2 in the genomic region around rs2842980 (100kb both upstream and downstream its genomic location). The plot also shows an intergenic genetic variant (rs2842992) associated to a higher risk of atrophic macular degeneration.

A similar trend was detected in the case of the rs9332 variant, located within the 3’UTR of the MTRR gene, encoding an enzyme essential for methionine synthesis (Supplementary Fig. 6A). This variant was reported to be associated with a higher risk of spina bifida, along with other variants within the same gene [44]. We found that the variant is associated with the increased relative expression of the short isoform of the MTRR transcript, as a consequence of the creation of a proximal canonical PAS. We can thus speculate that, similarly to what was shown for IRF5, this post-transcriptional event could lead to a variation in the activity of the enzyme activity and ultimately to increased disease susceptibility.

The same mechanism might provide putative mechanistic explanations for associations found by GWAS studies. For example we found the variant rs5855 to be an apaQTL for the PAM gene (Supplementary Fig. 6B), essential in the biosynthesis of peptide hormones and neurotransmitters [45, 46, 47]. No eQTLs or trQTLs for this gene were revealed by the analysis of the same data reported in [8]. This variant replaces an alternative PAS motif (AGUAAA) with the canonical AAUAAA, thus presumably increasing its strength. This PAS motif is located 26 bps upstream of an APA site corresponding to a 3’UTR of *∼*450 bps, instead of the *∼*2,000 bps of the canonical isoform, lacking several predicted microRNA binding sites. Indeed, our analysis revealed a shortening of the 3’UTR in individuals with the alternate allele, i.e. the canonical PAS motif. Notably, the variant is in strong LD (*R*^2^ = 0.90) with the intronic variant rs10463554, itself an apaQTL for PAM, which has been associated to Parkinson’s disease in a recent meta-analysis of GWAS studies [48].

The creation of a distal canonical PAS motif can lead instead to 3’ UTR lengthening, as shown by variant rs2842980 located in the 3’ UTR of SOD2 (Fig. 6C,D). The alternate allele creates a canonical AUUAAA PAS site, whereas the reference allele is UUUAAA, 19 bps upstream of the most distal annotated poly(A) site. Also this variant is in strong LD (*R*^2^ = 0.93) with an intergenic GWAS hit, namely variant rs2842992, which has been associated to atrophic macular degeneration [49]. A mechanistic involvement of SOD2 in macular degeneration is supported by a mouse model in which the gene was deleted in the retinal pigment epithelium, inducing oxidative stress and key features of age-related macular degeneration [50].

Conversely, the destruction of a canonical, proximal PAS motif leads to shortening of the 3’ UTR of BLOC1S2 (Supplementary Fig. 6C). The variant rs41290536 replaces the canonical PAS motif AAUAAA with the non-canonical one AAUGAA 17 bps upstream of a poly(A) site corresponding to a UTR length of *∼*750 bps compared to the *∼*2,200 of the longest isoform. The variant is in complete LD (*R*^2^ = 1) with two variants that have been associated to predisposition to squamous cell lung carcinoma (rs28372851 and rs12765052) [51].

#### 2.3.2 Alteration of microRNA binding

In an alternative scenario, genetic variants can influence the relative expression of alternative 3’UTR isoforms by acting on the stability of transcripts, for example through the creation or destruction of microRNA binding sites. For each gene with alternative 3’UTR isoforms, we divided the 3’ UTR into two segments: the “PRE” segment, common to both isoforms, and the “POST” segment expressed only by the longer isoform. Variants altering microRNA binding sites located in the POST segment can result in the variation of the relative isoform expression since they affect only the expression of the long isoform.

For example, we found that the rs8984 variant is associated with an increased prevalence of the long transcript isoform of the CHURC1 gene, an effect that could be due to the destruction of a binding site recognized by microRNAs of the miR-582-5p family within the POST segment of the gene (Supplementary Fig. 7). More generally, we found that apaQTLs are enriched, albeit slightly, among the genetic variants that create or break putative functional microRNA binding sites (OR = 1.15, 95% CI = 1.02 - 1.30, P-value = 0.022). However, we could not find significant agreement between the predicted and actual direction of the change in isoform ratios for these cases. Together with the marginal significance of the enrichment, this result suggests that the alteration of microRNA binding sites is not among the most relevant mechanisms in the genetic determination of 3’ UTR isoform ratios.

#### 2.3.3 Alteration of RNA-protein binding

RNA-binding proteins (RBPs) play important roles in the regulation of the whole cascade of RNA processing, including co- and post-transcriptional events. Although many of them have not been fully characterized yet, a collection of 193 positional weight matrices (PWMs) describing a large number of RNA motifs recognized by human RBPs has been obtained through in-vitro experiments [52]. Here we exploited this resource to identify SNPs that alter putative functional RBP binding sites. Consistently we the involvement of RBPs in the regulation of alternative polyadenylation, mRNA stability and microRNA action, we found a highly significant enrichment of RBP-altering SNPs among intragenic apaQTLs (OR = 1.48, 95% CI = 1.31 - 1.66, P-value = 8.47 *×* 10^*−*11^).

Specifically, we obtained a positive and significant OR for 20 individual RBP binding motifs (Supplementary Table 1). Although in most cases the enrichment is modest, some of the enriched motifs correspond to RNA-binding domains found in RBPs with a previously reported role in polyadenylation regulation (members of the muscleblind protein family [30, 53], KHDRBS1 [54] and HNRNPC [40]). Other enriched RNA-binding motifs are associated with splicing factors (RBM5, SRSF2, SRSF9 and RBMX) and other RBPs that may be involved in RNA processing (such as members of the MEX3 protein family and HNRNPL). On the contrary, only one significant motif is associated with a RBP that may be involved in RNA degradation (CNOT4 [55]). The involvement of several splicing factors is consistent with evidence supporting a mechanistic interplay between polyadenylation and splicing, that goes beyond the regulation of the usage of intronic poly(A) sites [56, 57, 58, 59, 60].

### 2.4 Extragenic apaQTLs act in-cis through the perturbation of regulatory elements

Understanding the function of extragenic apaQTLs is less straightforward because, although there are few examples of DNA regulatory elements contributing to APA regulation [14], it is commonly believed that APA is mainly controlled by cis-elements located within transcripts, both upstream and downstream of the poly(A) sites [13].

To further explore this aspect we took advantage of a different annotation of active genome regions, which includes the association between regulatory regions and target genes, namely the cis-regulatory domains (CRDs) identified in lymphoblastoid cell lines in Ref. [61]. Extragenic apaQTLs were indeed found to be enriched in CRDs (OR = 1.73, 95% CI = 1.69 - 1.78, P-value *<* 10^*−*16^). The 3D structure of the genome is a key aspect of gene regulation [62], as it determines physical contacts between distal regulatory regions and proximal promoters. In particular, CRDs have been described as active sub-domains within topologically associated domains (TADs), containing several non-coding regulatory elements, both proximal and distal. The perturbation of those regulatory elements by genetic variants can lead to the alteration of gene expression and perhaps interfere with other processes such as alternative polyadenylation, as suggested by our results. Importantly, CRDs have been assigned to the nearby genes they regulate. We could thus observe that extragenic apaQTLs tend to fall within CRDs that have been associated with their target genes much more frequently than expected by chance. Indeed, this correspondence was verified for 27,527 extragenic apaQTLs, while the same degree of concordance was never obtained in 100 permutations in which each extragenic apaQTL was randomly associated to a gene in its cis-regulatory window (median number of correspondences 12,571). These results suggest an important role of genetic variants located in active, non-transcribed cis-regulatory regions in regulating alternative polyadenylation of the target genes.

### 2.5 A role for apaQTLs in complex diseases

Since common genetic variation is involved in complex diseases, often by affecting gene regulation, a natural question is whether apaQTLs can be used to provide a mechanistic explanation for some of the genetically driven variability of complex traits, thus adding 3’ UTR length to the list of useful intermediate phenotypes. Besides the specific examples discussed above, we found an overall striking enrichment among apaQTLs of genetic variants reported in the NHGRI-EBI GWAS Catalog [63] (OR = 3.17, 95% CI = 3.01 - 3.34, P-value < 10^−16^).

We also investigated the enrichment of each trait category defined by the Experimental Factor Ontology (EFO) and then for each individual trait. In line with the fact that the apaQTL mapping was performed in lymphoblastoid cells, the strongest enrichment was observed for immune system disorders (OR = 5.41, 95% CI = 4.52 - 6.45, P-value = 2.50 *×* 10^*−*77^) (Fig. 7 and Supplementary Table 2). However, a strong enrichment was also detected for almost all the other tested categories, including neurological disorders (OR = 4.32, 95% CI = 3.86 - 4.83, P-value = 2.47 *×* 10^*−*142^) and cancer (OR = 3.96, 95% CI = 3.36 - 4.64, P-value = 4.15 *×* 10^*−*63^).

**Figure 7:**
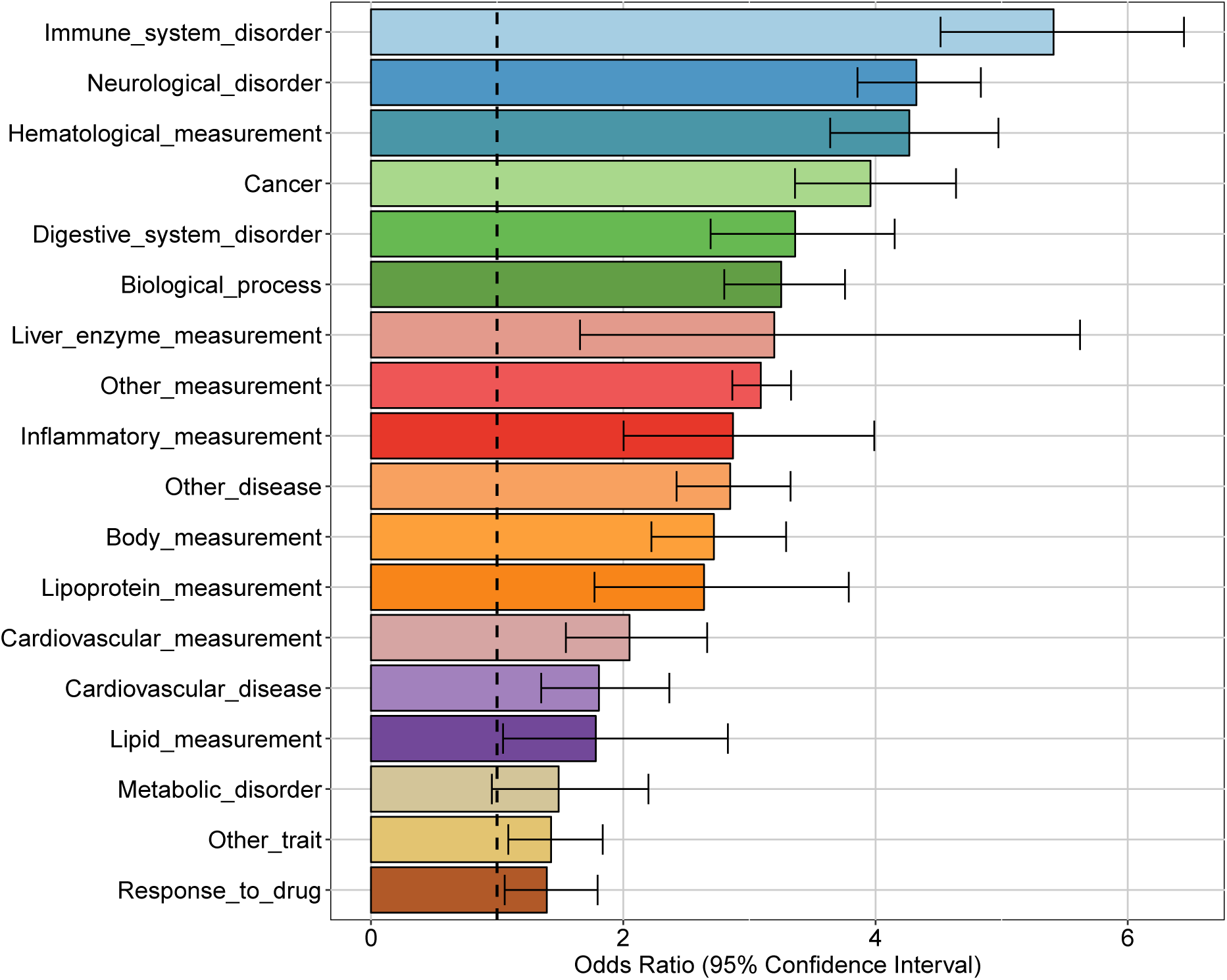
Enrichment of GWAS hits among apaQTLs, for different categories of complex traits. For each category, the OR obtained by logistic regression and its 95% CI are shown.

A significant enrichment was detected for 95 individual complex traits, including several diseases. Among these, the largest ORs were observed for autism spectrum disorder (OR = 42.6, 95% CI = 32.9 - 55.5, P-value = 2.36 *×* 10^*−*174^), squamous cell lung carcinoma (OR = 26.1, 95% CI = 15.7 - 43.3, P-value = 1.29 *×* 10^*−*36^), lung carcinoma (OR = 17.9, 95% CI = 12.7 - 25.2, P-value = 9.63 *×* 10^*−*62^), schizophrenia (OR = 10.6, 95% CI = 9.01 - 12.4, P-value = 1.25 *×* 10^*−*182^), and HIV-1 infection (OR = 6.51, 95% CI = 3.75 - 10.8, P-value = 2.28 *×* 10^*−*12^). The complete list of enriched traits can be found in Supplementary File 5.

We observed that apaQTLs that are also GWAS hits often map to genes in the human leukocyte antigen (HLA) locus, suggesting that in at least some cases the enrichment could be mostly driven by this genomic region. Somewhat unexpectedly, this was particularly evident for neurological disorders. In order to clarify this point, we evaluated all enrichments after excluding the variants in the HLA locus. Although in some cases the OR decreased after removing HLA variants, for most GWAS categories the enrichment was still significant (Supplementary Fig. 8 and Supplementary Table 3). For example, we found 155 apaQTLs associated with autism spectrum disorder, 116 of which affecting HLA genes. After the exclusion of HLA variants, the enrichment was still highly significant (OR = 10.66, 95% CI = 6.92 - 15.95, P-value = 7.05 *×* 10^*−*29^). On the contrary, the enrichment of variants associated to pulmonary adenocarcinoma is driven by the HLA locus, and becomes non-significant after excluding HLA variants (OR = 1.35, 95% CI = 0.22 - 4.39, P-value = 0.68). The complete list of enriched traits after the exclusion of HLA variants can be found in Supplementary File 6.

### 2.6 The effect of genetic variants on APA can be confirmed in patients

As briefly discussed above, the rs10954213 variant is associated with a higher risk of SLE. Evidence about the related molecular mechanism arose from the analysis of cell lines derived from healthy individuals [41, 42], and the effect of the variant on IRF5 expression in blood cells was confirmed in SLE patients [64, 65]. However, direct evidence on the effect of this variant on APA regulation in SLE patients is still missing.

In order to assess whether rs10954213 affects IRF5 APA regulation in SLE patients, we analyzed RNA-Seq data derived from whole blood cells in 99 patients [66]. We detected a strong difference in IRF5 *m/M* values among the three rs10954213 genotypes, with the alternative allele associated with higher *m/M* values, i.e. shorter 3’ UTR (Kruskal-Wallis test P-value = 9.21*×*10^*−*10^; Fig. 8). Therefore the variant has, at least qualitatively, the same effect in the whole blood of SLE patients as in lymphoblastoid cell lines of normal individuals.

**Figure 8:**
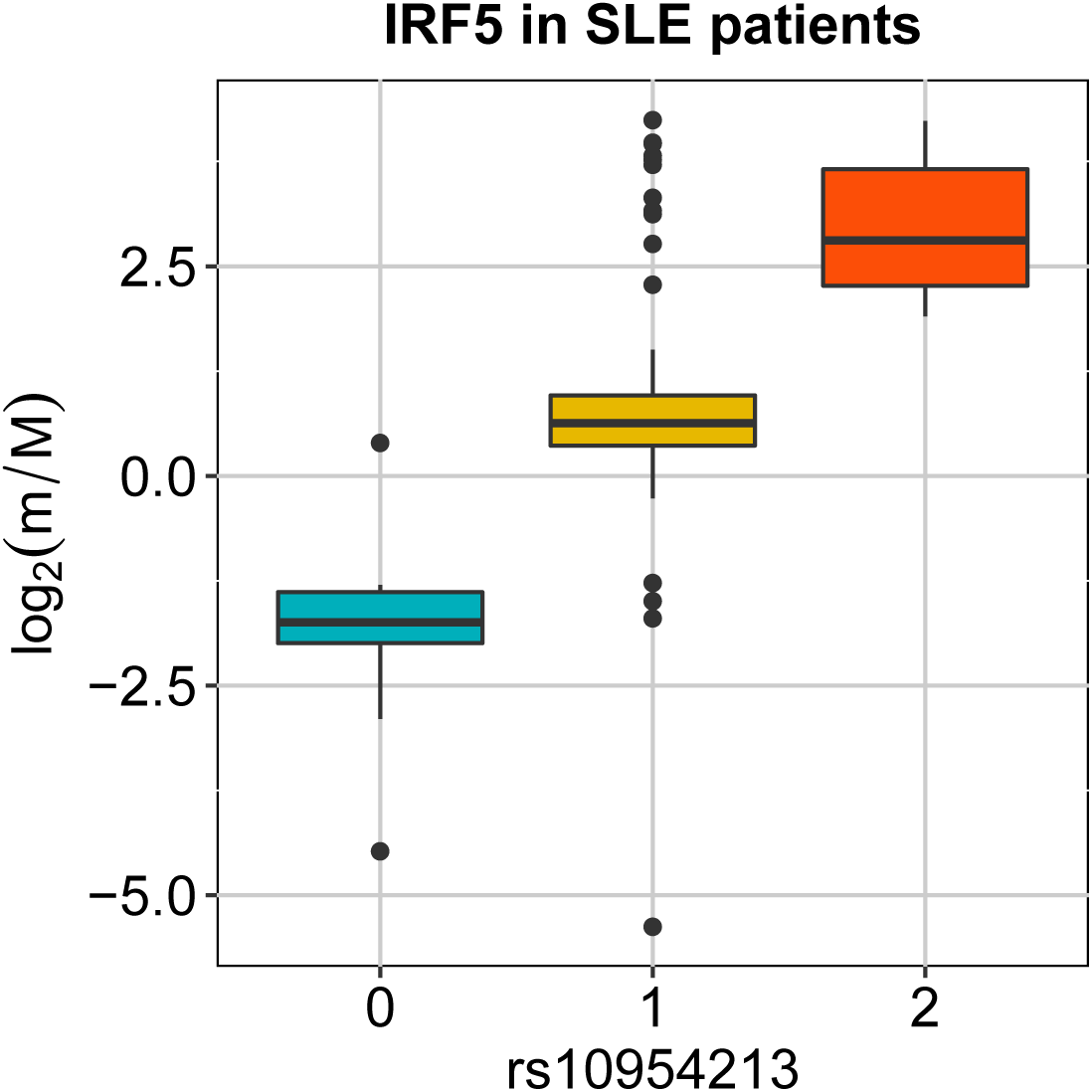
The effect of rs10954213 on the relative expression of the IRF5 alternative isoforms was investigated also in a small cohort of SLE patients. The boxplot show the variation of the log2-transformed m/M values obtained for IRF5 as a function of the genotype of the individuals.

## 3 Discussion

We used a new efficient strategy to study how human genetic variants influence the expression of alternative 3’UTR isoforms. This issue has been previously investigated with different approaches [26, 25, 27, 8, 9]. The method we propose combines wide applicability, being based on standard RNA-Seq data, with the high sensitivity allowed by limiting the analysis to a single type of transcript structure variant, namely 3’ UTR length. Such higher sensitivity led us to discover thousands of variants associated with 3’ UTR length that were not identified in a general analysis of trascripit structure from the same data in [8]. Moreover, the significant overlap between our apaQTLs and the eQTLs identified in [8] confirms the known relevant role of 3’UTRs in gene expression regulation. However, the regulation of 3’ UTR length is known to affect regulatory processes that do not directly alter mRNA abundance, such as regulation of translation efficiency, mRNA localization and membrane protein localization [12, 36]. Indeed most of the apaQTLs we found were not identified as eQTLs in [8].

The various mechanisms underlying the association between genetic variants and the relative abundance of 3’UTR isoforms can be classified in two main classes based on whether they affect the production or degradation rates of the isoforms. The production related-mechanisms include the alteration of APA sites, of cis-regulatory elements located in promoters and enhancers, and of binding sites of RBPs involved in nuclear RNA processing; the degradation-related mechanisms include the alteration of the binding sites of microRNAs and cytoplasmatic RBPs affecting mRNA stability. Taken together, our results suggest that the genetic effects on 3’UTR isoforms act prevalently at the level of production, as shown by the strong enrichment of apaQTLs in non-transcribed regulatory regions and among the variants creating or disrupting APA sites, and by the relatively weak enrichment of variants creating or disrupting microRNA binding sites. Also the results on altered RBP binding sites confirm this picture, since most motifs altered by apaQTLs are associated to nuclear RBPs involved in nuclear RNA processing.

In particular, we identified several apaQTLs creating or destroying putative functional PAS motifs. However, it should be noted that our ability to detect these events is intrinsically limited by the motif repertoire that we used [40], which might miss some of the rarest alternative PAS motifs. For example, we found that the rs6151429 variant is associated with the increased expression of the long isoform of the transcript codified by the Arylsulfatase A (ARSA) gene (Supplementary Fig. 6D), in agreement with previous evidence [67]. However, we did not include this variant among those disrupting a PAS motif since the disrupted motif (AAUAAC) is not included in the catalog that we used. In addition, we considered only PAS-altering single nucleotide substitutions, while also other types of genetic variants can modify the PAS landscape of a gene. For example, a small deletion (rs374039502) causes the appearance of a new PAS motif within the TNFSF13B gene, and has been associated with an higher risk of both multiple sclerosis and SLE in the Sardinian population [68].

We observed a strong enrichment of apaQTLs in regulatory regions such as promoters and enhancers, as previously found for variants generically affecting transcript structure in [8]. These results point to an important role of DNA-binding cis-acting factors in the regulation of 3’ UTR length, and to the existence of a widespread coupling between transcription and polyadenylation [12, 69]. Moreover, it has been shown that RBPs involved in APA regulation can interact with promoters [14].

Regarding the effect of genetic variants on mRNA stability, we focused on the perturbation of microRNA binding, taking into account both the creation and the destruction of microRNA binding sites within transcripts. The relevance of mRNA stability seemed to be confirmed by a modest enrichment of microRNA-altering SNPs among intragenic apaQTL, however the direction of their effect on microRNA binding is not statistically consistent with the expected direction of the change in 3’ UTR isoform ratio. The same type of ambiguity has been previously reported with regard to the relationship between the effect of SNPs on microRNA binding and gene expression levels [70] and makes us doubt whether these microRNA-altering apaQTLs are truly causal for the associated gene. These results suggest that the alteration of microRNA binding may not be a predominant mechanism explaining the variation of the expression of alternative 3’UTR isoforms across individuals. Limitations in the accuracy of predicted micorRNA binding sites might also contribute to this result.

Another possible mechanism of action of intragenic apaQTLs is the perturbation of the regulatory action of RBPs, as indicated by the modest but highly significant enrichment of SNPs altering RNA-binding motifs. However, the lack of strong enrichments when considering each motif individually suggests that specific RBP motifs may have a small regulatory impact on APA that may also depend on the context, as recently suggested [30]. As in the case of microRNAs, also our limited knowledge of the binding preferences of RBPs might limit our power to detect their effects: More sophisticated models should take into account the highly modular structure of RBPs that often include multiple RNA binding domains (RBDs), the emerging importance of both the binding context and the RNA structure and even more sophisticated modes of RNA binding [71, 72].

Furthermore, it is reasonable to assume that also non-canonical modes of APA regulation can be affected by genetic variants and therefore drive the detection of variable isoform expression ratios. For example, it has been recently suggested that an epitranscriptomic event, the m^6^A mRNA methylation, can be associated with alternative polyadenylation [15]. In addition, recently published results suggest that genetic variants could affect APA regulation also in an indirect way, without affecting the regulatory machinery. Past studies have reported that a narrow range of 10-30nt between the PAS and the poly(A) site is required for efficient processing, however [73] suggested that also greater distances can sometimes be used thanks to RNA folding events that bring the PAS and the poly(A) site closer to each other. Therefore, we can speculate that if a genetic variant affects RNA folding in such a way as to modify the distance between the PAS and the poly(A) site, it could also influence APA regulation.

While the mechanisms discussed above act at the level of the primary or mature transcript, our results revealed a perhaps unexpectedly large number of extragenic apaQTLs, mostly located in regulatory regions. These apaQTLs point to an important role of DNA-binding elements such as transcription factors in regulating alternative polyadenylation through long-distance interactions with cleavage and polyadenylation factors. The investigation of these mechanisms is thus a promising avenue of future research.

Alternative polyadenylation can affect several biological processes, influencing mRNA stability, translation efficiency and mRNA localization [13]. Therefore, it is not surprising that its perturbation has been associated with multiple pathological conditions [7, 22]. In the present study, we detected a strong enrichment of GWAS hits among apaQTLs, supporting the idea that 3’ UTR length is a useful addition to the list of intermediate molecular phenotypes that can be used for a mechanistic interpretation of GWAS hits. In particular, we identified genetic variants previously associated to neurological disorders, such as autism, schizophrenia and multiple sclerosis, which may act by affecting the regulation of polyadenylation. The importance of post-transcriptional events in the onset of neurological diseases has been recently confirmed by two studies demonstrating that genetic variants affecting alternative splicing (sQTL) give a substantial contribution to the pathogenesis of schizophrenia [74] and Alzheimer’s disease [75]. We also observed that the relevant apaQTLs often map to HLA genes, but that the enrichment is not explained by the HLA locus alone. On the other hand, examples of APA events involving HLA genes have been reported [76, 77] and genes encoding antigen-presenting molecules account for the highest fraction of genetic risk for many neurological diseases [78].

A gene-based alternative approach to the interpretation of GWAS has been recently proposed. In the original implementation of Transcriptome Wide Association Studies (TWAS) [79], eQTL data obtained in a reference dataset are used to predict the genetic component of gene expression in GWAS cases and controls, which is then correlated with the trait of interest, thus allowing the identification of susceptibility genes. More recently, the authors of [80] proposed a summary-based TWAS strategy in which the association between the genetic component of gene expression and a trait is indirectly estimated through the integration of SNP-expression, SNP-trait, and SNP-SNP correlation data. Furthermore, this kind of analysis has also been performed exploiting a collection of sQTLs, leading to the identification of new susceptibility genes for schizophrenia [81] and Alzheimer’s disease [75]. In a similar way, apaQTLs could be used to discover cases in which the association between genes and diseases is driven by the alteration of the expression of alternative 3’UTR isoforms.

We are aware of some limitations of this study. First, the simple model that we used for the definition of alternative 3’UTRs isoforms limits the type of events that can be detected, because we can see only events involving poly(A) sites located within the transcript segments taken into account for the computation of the *m/M* values (the PRE and the POST segments). Nonetheless, the adoption of this simple model significantly reduces the computational burden and might be sufficient to indicate general trends that can be subsequently further investigated with more sophisticated models. Indeed, it has been previously shown, in a slightly different context (i.e. the comparison of APA events detected in different cellular conditions or tissues), that the results obtained with our model are comparable with those obtained exploiting a more complex model that takes into account all the possible APA isoforms of a gene, especially because also genes with multiple poly(A) sites mainly use only two or a few of them [29]. Second, our strategy depends on a pre-existing annotation of poly(A) sites: Methods that infer the location of poly(A) sites from RNA-Seq data are available, but they can have lower sensitivity in the detection of APA events [29, 30]. In addition, we examined only a single cell type (lymphoblastoid cells) to demonstrate the feasibility of apaQTL mapping analysis. A broader investigation, exploiting data such as those provided by Genotype-Tissue Expression (GTEx) consortium [82], would be particularly valuable. Indeed, APA regulation seems to be significantly tissue-specific and global trends of poly(A) sites selection in specific human tissues have been described: For example transcripts in the nervous system and brain are characterized by preferential usage of distal PAS, whereas in the placenta, ovaries and blood the usage of proximal PAS is preferred [12].

In conclusion, we have identified thousands of common genetic variants associated with alternative polyadenylation in a population of healthy human subjects. Alternative polyadenylation is a promising intermediate molecular phenotype for the mechanisitic interpretation of genetic variants associated to phenotypic traits and diseases.

## 4 Material and Methods

### 4.1 Data sources

#### 4.1.1 Human genome and transcriptome

The coordinates of the NCBI Reference Sequences (RefSeqs) in the human genome (hg19) were downloaded from the UCSC Genome Browser (09/04/2015) [83, 84]. The corresponding transcript-gene map was downloaded from NCBI (version 69) and the Bioconductor R package `org.Hs.eg.db v3.4.0` [85] was used to associate each Entrez Gene Id to its gene symbol. In addition, the reference sequence of the hg19 version of the human genome was downloaded from the ENSEMBL database and a collection of poly(A) sites was obtained from PolyA_DB2 (10/02/2014) [31].

ChromHMM annotations [37] were downloaded from the UCSC Genome Browser for the GM12878 and the NHEK cell lines (http://genome-euro.ucsc.edu/cgi-bin/hgFileUi?db=hg19&g=wgEncodeBroadHmm) In addition, the coordinates of Cis Regulatory Domains (CRDs) and their association with genes were downloaded for lymphoblastoid cells from ftp://jungle.unige.ch/SGX/ [61].

### 4.1.2 WGS and RNA-Seq data

We exploited the RNA-Seq data obtained by the GEUVADIS consortium in lymphoblastoid cell lines of 462 individuals belonging to different populations, but we considered only 373 individuals with European ancestry (EUR). BAM files were downloaded from the E-GEUV-1 dataset [8] in the EBI ArrayExpress archive (https://www.ebi.ac.uk/arrayexpress/files/E-GEUV-1/processed). We also downloaded genotypic data for the same individuals (https://www.ebi.ac.uk/arrayexpress/files/E-GEUV-1/genotypes/) and the results of the eQTL/trQTL map-ping analyses (https://www.ebi.ac.uk/arrayexpress/experiments/E-GEUV-1/files/analysis_results/). The downloaded VCF files include genotypes for 465 individuals: Among the 462 of them for which also RNA-Seq data are available, the large majority had been previously subjected to Whole Genome Sequencing (WGS) by the 1000 Genome Project (Phase 1) [3], but the GEUVADIS consortium additionally obtained genomic data for 41 of them through genotyping with Single Nucleotide Polymorphism (SNP) array followed by genotype imputation [8]. Furthermore, whole blood RNA-Seq data for 99 individuals affected by SLE were downloaded from the NCBI SRA database (SRP062966) [86, 66].

#### 4.1.3 Regulatory motifs and related expression data

Different collections of regulatory motifs were downloaded. A list of 18 PAS motifs was obtained from [40], microRNA seeds were downloaded from TargetScan 7.2 [87] and Positional Weight Matrices (PWMs) describing the binding specificities of RNA-binding proteins were downloaded from the CISBP-RNA dataset [52], including both the experimentally determined motifs and those that were inferred from related proteins. In addition, the list of microRNAs and RBPs expressed in lymphoblastoid cells were obtained from the expression data available in the E-GEUV-2 dataset [8] on the EBI ArrayExpress archive (https://www.ebi.ac.uk/arrayexpress/files/E-GEUV2/GD452.MirnaQuantCount.1.2N.50FN.samplename.resk10.txt and https://www.ebi.ac.uk/arrayexpress/files/E-GEUV-1/GD462.GeneQuantRPKM.50FN.samplename.resk10.txt.gz).

#### 4.1.4 GWAS Catalog

A collection of genomic loci associated with human complex traits was obtained by downloading the NHGRI-EBI GWAS Catalog, v1.0.2 [63]. This resource is continuously updated: the version we used was downloaded on October 10^th^, 2018 and it was mapped to GRCh38.p12 and dbSNP Build 151. From the same website, we also downloaded a file showing the mapping of all the reported traits to the Experimental Factor Ontology (EFO) terms [88], including the parent category of each trait (the version of the downloaded file was r2018-09-30). In addition, the dpSNP Build 151 [89] collection of human genetic variants was downloaded for hg19.

### 4.2 Annotation of alternative 3’UTR isoforms

We considered the human transcripts included in RefSeq and associated them with the corresponding Entrez Gene Id. Moreover, we collapsed together the structures of all the transcripts assigned to a gene, using the union of all the exons of the various transcripts associated to a gene and defining the 3’ or 5’ UTR using respectively the most distal coding end and the most proximal coding start. The annotation of the resulting gene structures can be found in the supplementary data (see Supplementary File 1).

The coordinates of the human poly(A) sites were converted from hg17 to hg19 using liftOver [90] and then combined with the gene structures defined above to define the alternative 3’UTR isoforms. For the definition of alternative 3’UTR isoforms we adopted a simple model taking into account only two alternative poly(A) sites for each gene, because previous evidence suggests that also genes with multiple poly(A) sites mainly use only two of them [29]. In particular, for each gene we selected the most proximal poly(A) site among those falling within exons, preferring those located within the 3’UTR, and the end of the gene as the distal poly(A) site. In this way we were able to define two segments of interest for each gene: the PRE segment, extending from the beginning of the last exon to the proximal poly(A) site, and the POST segment, from the proximal poly(A) site to the end of the gene. The PRE fragment is assumed to be expressed by both the long and the short isoform, while the POST segment should be expressed exclusively by the long isoform. The GTF file used for the computation of *m/M* values is available as Supplementary File 2.

The relative prevalence of the short and long isoforms are evaluated, as described below, based on the number of RNA-Seq reads falling into the PRE and POST regions. While the whole region from the transcription start site to the proximal poly(A) site could be taken, in principle, as the PRE region, we chose to limit it to the last exon to minimize the confounding effect of alternative splicing.

### 4.3 Computation of m/M values

Using the Bioconductor R package Roar [29], for each gene with alternative 3’UTR isoforms we obtained an *m/M* value in each individual. The *m/M* value estimates the ratio between the expression of the short and the long isoform of a gene in a particular condition and the *m/M* _a,i_ of gene *a* in the *i_th_* individual is defined as

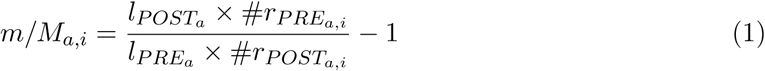

where *l*_*PRE*_*a*__ and *l*_*POST*_*a*__ are respectively the length of the PRE and POST segment of the gene *a*, #*r*_*PRE*_*a,i*__ and #*r*_*POST*_*a,i*__ are respectively the number of reads mapped on the PRE and the POST segment of the gene *a* in the *i*_*th*_ individual.

The *m/M* values were computed for 14,542 genes for which we were able to define alternative 3’UTR isoforms. Infinite and negative values of *m/M* (that happen when the POST region does not produce any reads, and when the POST region produces more reads than the PRE region after length normalization, respectively) were considered as missing values. Then only those on autosomal chromosomes (chr1-22) and with less than 100 missing *m/M* values were selected for the following investigation, leaving us with 6,256 genes.

### 4.4 Genotypic data pre-processing

Starting from the downloaded VCF files, we extracted genotypic data for 373 EUR individuals for whom also RNA-Seq data are available using `VCFtools` [91]. In addition, only common genetic variants with Minor Allele Frequency (MAF) higher than 5% were considered in all the following analyses; the MAF value of each genetic variant was reported in the VCF files, but we recomputed them taking into account that the reference allele in the VCF file may not always be the most frequent one in the EUR population considered by itself, and conservatively attributing the most frequent homozygous genotype to individuals for which the genotype was missing.

### 4.5 Principal Component Analysis of genotypic data

It is known that special patterns of linkage disequilibrium (LD) can cause artifacts when a Principal Component Analysis (PCA) is used to investigate population structure [92]. We filtered out all the genetic variants falling within 24 long-range LD (LRLD) regions whose coordinates were derived from [92]. In addition, following [93], we performed an LD-pruning of the genetic variants using the `--indep-pairwise` function from `PLINK v1.9` [94] to recursively exclude genetic variants with pairwise genotypic R^2^ *>*80% within sliding windows of 50 SNPs (with a 5-SNPs increment between windows). Also in this case `VCFtools` [91] was used to apply all these filters to the VCF files and finally `EIGENSTRAT v6.1.4` [95] was used to run the PCA on the remaining genotypic data at the genome-wide level.

### 4.6 apaQTL mapping

From a statistical point of view, we adopted the same strategy used in standard eQTL mapping analyses [8] to identify genetic variants that influence the expression level of the alternative 3’UTR isoforms of a gene. For each of the 6,256 examined genes, we defined a cis-window as the region spanning the gene body and 1 Mbp from both its TSS and its TSE. Then, for each gene a linear model was fitted, independently for each genetic variant within its cis-window, using the genotype for the genetic variant as the independent variable and the log2-transformed *m/M* value of the gene as the dependent variable:

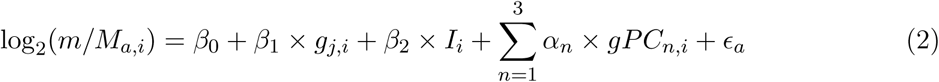

where *log*_2_(*m/M_a,i_*) is the log2-transformed *m/M* value computed for the *a* gene in the *i_th_* individual, *g_v,i_* is the genotype of the *i_th_* individual for the *j_th_* genetic variant, *I_i_* is the imputation status (0|1) of the *i_th_* individual, *gPC_n,i_* is the value of the *n_th_* Principal Component (PC) obtained from genotypic data for the *i_th_* individual, *β*_0_ is the intercept, *β*_1_, *β*_2_ and *α_n_* are the fitted regression coefficients and *ϵ_a_* is the error term for the gene *a*.

The fitting of the linear models was done using the CRAN R package `MatrixEQTL` [96]. Genotypes were represented using the standard 0/1/2 codification, referring to the number of alternative alleles present in each individual, and matrices with genotypic information were obtained from VCF files exploiting the Perl API (Vcf.pm) included in the VCFtools suite [91]. Following [8], in all our models we included both the imputation status of the individuals and the first three PCs obtained from genotypic data as covariates, in order to correct for possible biases due to population stratification (Supplementary Fig. 1) or genotype imputation.

The observed distribution of nominal P-values was compared with the expected one in Quantile-Quantile plots (Q-Q plots), revealing the expected inflation due to the LD issue (Supplementary Fig. 2). A permutation-based procedure was implemented [97]: all the models were fitted again after the random shuffling of the *m/M* values of each gene across samples; then for each gene-variant pair we counted how many times we obtained a random P-value less than its nominal P-value and divided this value by the total number of random tests done. Finally, to control for multiple testing, the empirical P-values were corrected with the Benjamini-Hochberg procedure [98] and models with a corrected empirical P-value less than 0.05 were considered statistically significant. Manhattan plots were drawn using the CRAN R package `qqman` [99].

### 4.7 Comparison with other molecular QTLs

In order to compare the genes for which we detected one or more apaQTLs with those for which eQTL/trQTL were reported [8], we translated the Ensembl Gene IDs (ENSG) to NCBI Entrez Gene IDs using Ensembl v67 [100] retrieved using the Bioconductor R package `biomaRt v2.30` [101, 102]. 229 ENSGs could not be translated with this procedure and were therefore excluded from this analysis.

### 4.8 Enrichment analyses

In order to functionally characterize the apaQTLs, we analyzed the enrichment of several features among such variants, including their genomic location, their ability to alter known regulatory motifs, and their association with complex diseases. All enrichments were evaluated through multivariate logistic regression to allow correcting for covariates. In this section we provide an overview of the method, but refer to the following subsections for details about each analysis.

For each feature we first established which genetic variants were potentially associated with the feature (for example only variants in the 3’ UTR can alter microRNA binding sites). Therefore, each enrichment analysis started with the selection of the “candidate variants” that were subsequently subjected to an LD-based pruning, in order to obtain a subset of independent candidate variants (the same strategy was implemented for example in [103] to evaluate the enrichment of GWAS hits among eQTLs). LD-based pruning was always performed using PLINK with the same parameters used in the case of the PCA of genotypic data (see above), but applied in each case to the candidate variants only. To each candidate variant surviving pruning we attributed a binary variable indicating whether it has the feature under investigation. Finally, these variants are classified as apaQTLs (i.e. corrected empirical P-value < 0.05 for at least one gene) and null variants (i.e. nominal P-value > 0.1 in all the fitted models). We excluded the “grey area” variants with nominal P-value < 0.1 but empirical corrected P-value > 0.05 as they are likely to contain many false negatives. Finally we fitted a multivariate logistic model in which the dependent variable is the apaQTL/null status of the variant, and the independent variables are the feature of interest and covariates. The latter always include the MAF of the variant, since variants with higher MAF are more likely to be found as significant apaQTLs, and possibly other covariates depending on the feature under examination (see below).

The logistic model can thus be written as:

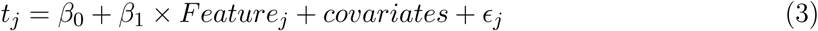

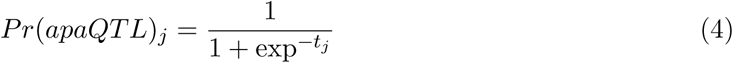

where *Feature_j_* is a binary variable indicating whether the genetic variant *j* has the feature of interest, *β*_0_ is the intercept, *β*_1_ is the regression coefficient for the feature, *∊_j_* is the error term and *Pr*(*apaQTL*)_*j*_ is the fitted probability that the genetic variant *j* is an apaQTL. As expected, in our models the regression coefficient of the MAF was always positive. The regression coefficient of the *Feature* term and its associated P-value were used to establish if having the feature under investigation influences the probability of being an apaQTL, and to compute the corresponding odds ratio (OR).

#### 4.8.1 Chromatin states

This analysis was performed independently for two cell types (the GM12878 and NHEK cell lines). In both cases, the candidate variants were virtually all the genetic variants for which the apaQTL models were fitted, but we excluded those not associated with any chromatin state and all the structural variants, because their length can prevent them from being univocally associated with a chromatin state.

Each of the 15 chromatin states and 6 broad chromatin classes (promoter, enhancer, insulator, transcribed, repressed and inactive) defined in [37], separately for the two cell lines, was treated as a binary feature to be used as a regressor in Eq. (3), with value 1 assigned to the variants falling within a DNA region associated to the given chromatin state. Only the MAF was included in the covariates.

#### 4.8.2 Gene regions

The candidate variants were all the intragenic variants for which the apaQTL models were fitted. We defined as intragenic all variants falling between the start and the end of the gene, plus 1,000 bps after the end (to take into account possible misannotations of the 3’ UTR).

Independent enrichment analyses were performed for the following sequence classes: coding exons, introns, 5’UTR and 3’UTR. For each class the binary feature used as a regressor was assigned the value 1 for variants falling within the class and 0 otherwise. Only the MAF was included in the covariates.

#### 4.8.3 Cis Regulatory Domains

The candidate variants were all the extragenic variants (i.e. all variants that are not intragenic according to the definition given above) for which an apaQTL model was fitted. The binary feature was given value 1 for variants falling within a CRD and 0 otherwise. Besides the MAF, the distance from the nearest gene was included as a covariate, since variants closer to a gene are more likely to be apaQTLs.

To verify that the apaQTLs tend to be included in the CRDs specifically associated to the gene on which they act, we translated the CRD-gene associations provided in [61] into Entrez Gene IDs, and we counted how many genetic variants fall within a CRD associated to at least one gene for which the variant is an apaQTL. This number was then compared with that obtained in the same way after randomly assigning a target gene to each extragenic variant within the cis-window used for apaQTL analysis (100 independent randomizations were used).

#### 4.8.4 Alteration of putative functional motifs

Similar strategies were implemented to investigate the alteration of different types of putative functional motifs by intragenic variants. This analysis was restricted to Single Nucleotide Polymorphism (SNPs), excluding therefore both indels and structural variants. For all SNPs we reconstructed the sequence of both the reference (REF) and the alternative (ALT) allele in the 20 bp region around each candidate genetic variant to determine whether the ALT allele creates or destroys a functional motif with respect to the REF allele. The functional motifs analyzed included PAS motifs, microRNA binding sites, and RBP binding sites.

To each candidate variant surviving LD pruning we associated, using PLINK, a list of tagging variants with genotypic R^2^ *>* 80%, and a binary feature value of 1 if the candidate variant itself or any of its tagging variant altered a functional motif. The enrichment of apaQTLs among motif-affecting variants was then evaluated with the logistic model described by Eq. 3. In the following, we describe the details of the logistic model for each class of functional motifs.

##### PAS motifs

The PAS motif is always located upstream of its target poly(A) site. It has been suggested that a narrow range of 10-30 nt is required for efficient processing, but recent work suggests that also larger distances can be functional thanks to RNA folding processes bringing the poly(A) site closer to the PAS [73]. Assuming that a PAS-altering SNP would affect the usage of its nearest poly(A) site, we associated to each intragenic SNP the nearest downstream poly(A) site, selected those for which such poly(A) site was located within the PRE/POST segments, and retained as candidate variants only those whose distance from the corresponding poly(A) site was between 10 and 100 nt. PAS-altering variants were defined as those for which a particular PAS motif was found in either the REF or the ALT sequence, but not in both (note that the interconversion between PAS motifs is considered as well, assuming that they can have different strength).

##### microRNA binding sites

microRNA binding sites located downstream of a poly(A) site, and hence in the POST segment, can affect the relative abundance of the long and short isoforms by allowing the selective degradation of the former by microRNAs. Therefore, we chose as candidate variants all the SNPs within the POST segment of the genes analyzed. Putative microRNA binding sites were classified, as in [87], in three classes: 8mer, 7mer-m8, and 7mer-A1 (matches classified as 6-mer were not considered). A variant was defined to alter a microRNA binding site if a putative binding site was present in either the REF or the ALT sequence, but not in both, or if the site class was different between the REF and the ALT sequences. Moreover, altering variants were classified as creating (destroying) a binding site if only the ALT (REF) sequence contained a binding site or if the ALT (REF) sequence contained a stronger binding site than the REF (ALT), according to the hierarchy 8mer > 7mer-m8 > 7mer-A1 match. Only microRNA families conserved across mammals or broadly conserved across vertebrates and expressed in lymphoblastoid cells were considered. Following [8], each microRNA was considered expressed if its expression value was greater than 0 in at least 50% of the samples, and each microRNA family was considered expressed if at least one of its microRNAs was expressed.

##### RBP motifs

The candidate variants were all the intragenic SNPs. FIMO [104] was used to scan the REF and ALT sequences around each candidate variant, using as background the nucleotide frequencies on the sequence of all the analyzed genes. A motif was considered altered if its score was greater than 80% the score of the perfect match in only one of two alleles. As in the case of microRNAs, only motifs corresponding to RBPs expressed in lymphoblastoid cell lines were considered. Enrichment was evaluated both for SNPs altering any RBP motif, and for each expressed RBP separately.

#### 4.8.5 GWAS hits

We considered only the GWAS catalog records referring to a single genetic variant on autosomal chromosomes for which all the fields CHR_ID, CHR_POS, SNPS, MERGED, SNP_ID_CURRENT, and MAPPED_TRAIT_URI were available, as well as the RSID. The coordinates of the selected genetic variants in hg19 were derived from dbSNP Build 151. We thus obtained 56,672 genetic variants associated with at least one complex trait. Furthermore, starting from the EFO URI(s) reported for each association, we obtained the corresponding EFO Parent URI(s) from the EFO annotation file.

All variants examined as potential apaQTLs were considered as our candidate variants. A binary feature value of 1 was attributed to each candidate variant surviving LD pruning and associated to a trait, or with a tagging variant associated to a trait, as in the case of motif-altering variants. Enrichment was evaluated for all trait-associated variants together, for each single trait, and for trait categories defined based on the EFO ontology. Only traits and trait categories associated with at least 100 GWAS hits were analyzed. The same analysis was also performed after excluding all variants within the HLA locus, as defined by The Genome Reference Consortium (https://www.ncbi.nlm.nih.gov/grc/human/regions/MHC?asm=GRCh370).

### 4.9 The rs10954213 variant in SLE patients

Since we were interested in the IRF5 gene only, RNA-Seq reads were aligned to a reduced genome comprising the gene sequence and an additional 50bp at its 3’ end using Bowtie v2.2.3 [105] and TopHat v2.0.12 [106]. As genotypic data were not available for these individuals, we inferred the rs10954213 variant status from the relative proportion of A and G in the RNA-Seq reads. Individuals were considered homozygous for the reference (G) or for the alternative (A) allele when the same nucleotide was present in all the reads, and a single read with a different nucleotide was considered sufficient to call an heterozygous individual (Supplementary Fig. 9). In this way we obtained 10 homozygotes for the reference allele, 73 heterozygotes and 16 homozygotes for the alternative allele. The MAF thus obtained is consistent with that reported in the NCBI dpSNP database [89]. A Kruskal-Wallis test was then used to evaluate the differences in *m/M* values between genotypes. A different criterion for the assignment of genotypes (at least 20% of the reads carrying the less frequent allele required to call a heterozygous individual) gave comparable results (Supplementary Fig. 10).

## Supplementary figures

**Supplementary figure 1:**
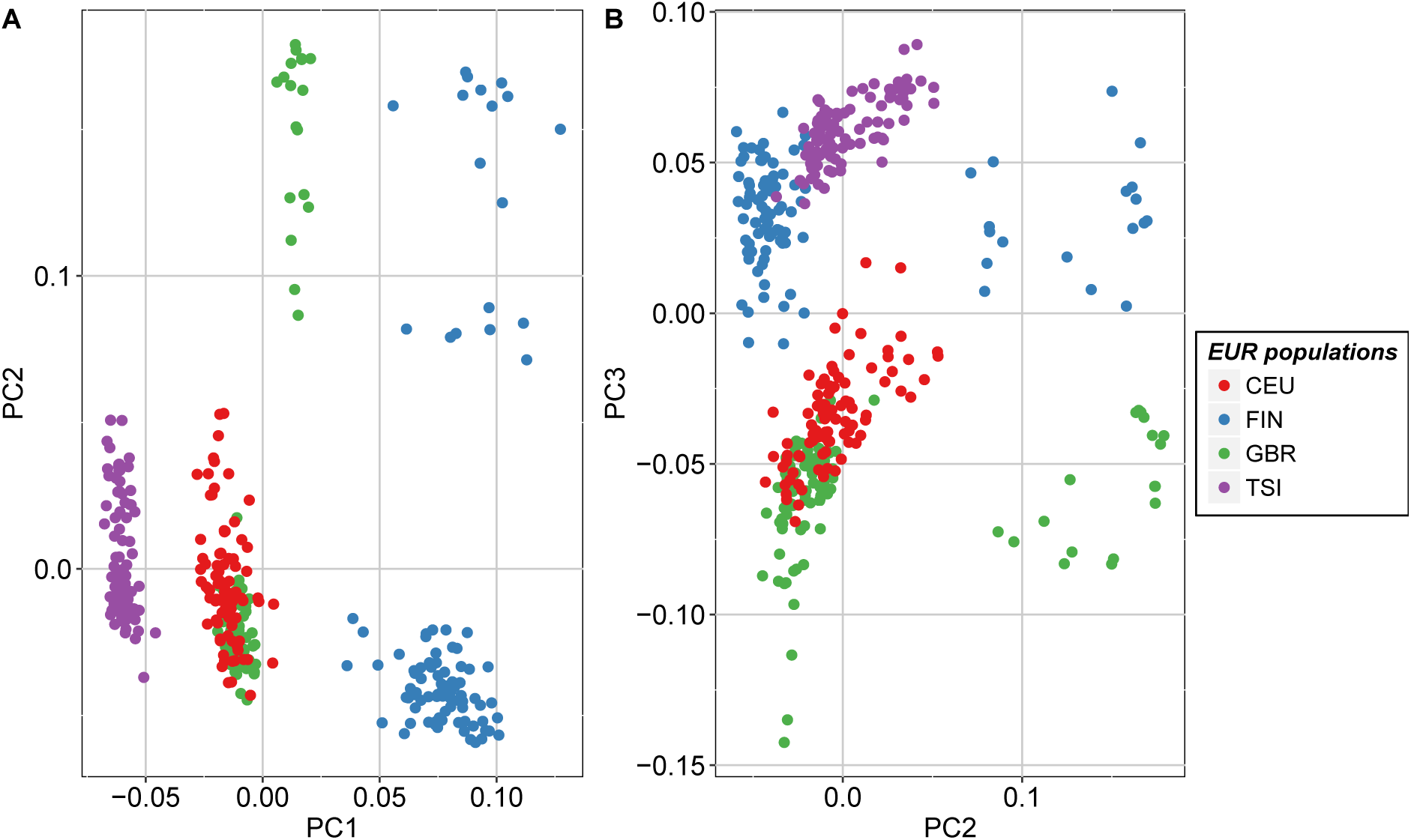
Principal Component Analysis (PCA) of the genotypic data of the EUR individuals were plotted. Points are colored according to the subpopulation of origin: Utah Residents (CEPH) with Northern and Western European Ancestry (CEU), Finnish in Finland (FIN), British in England and Scotland (GBR) and Toscani in Italia (TSI).

**Supplementary figure 2:**
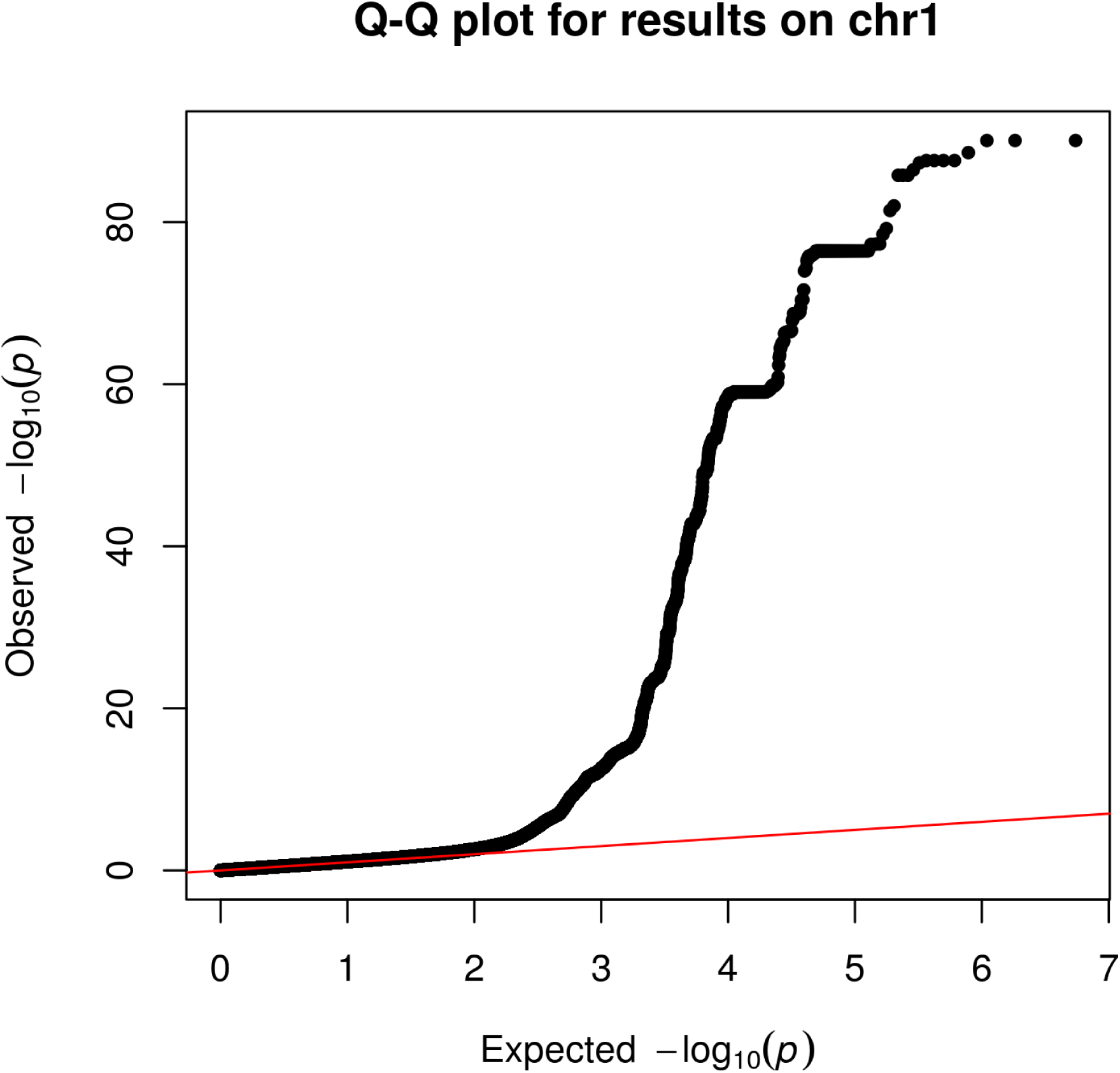
Q-Q plot comparing the distribution of P-values obtained fitting apaQTL models for genes on chr1 with the expected uniform distribution. It was generated by the CRAN R package qqman.

**Supplementary figure 3:**
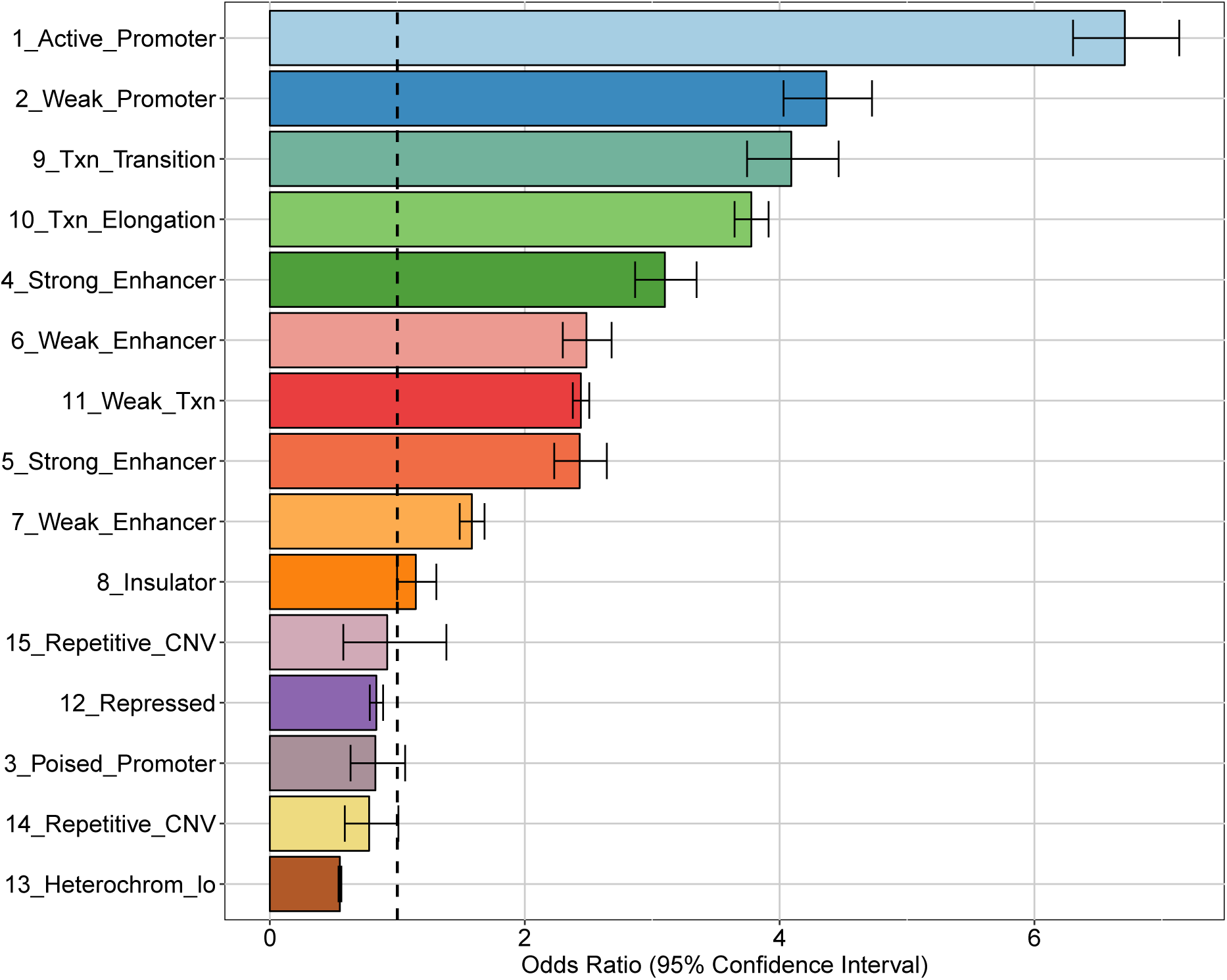
Enrichment of apaQTLs within chromatin states, taking into account all the 15 chromatin states reported in the ChromHMM annotation. For each of them, the OR obtained by logistic regression and its 95% CI are shown.

**Supplementary figure 4:**
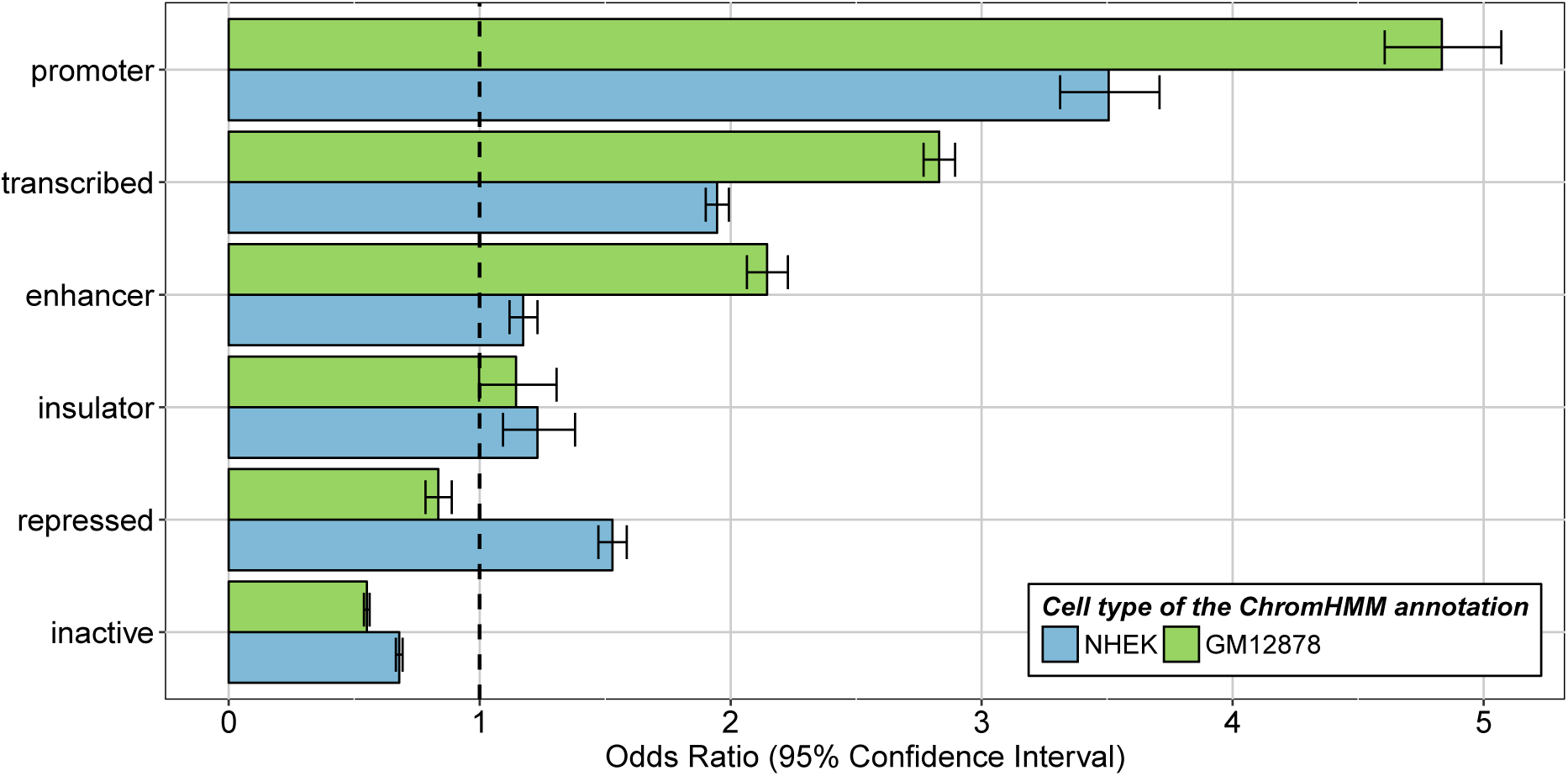
The results of the enrichment analysis performed with the broad chromatin states of the relevant cell type were compared with those obtained using the ChromHMM annotation of another cell type (NHEK). For each category, the OR obtained by logistic regression and the corresponding 95% CI are shown.

**Supplementary figure 5:**
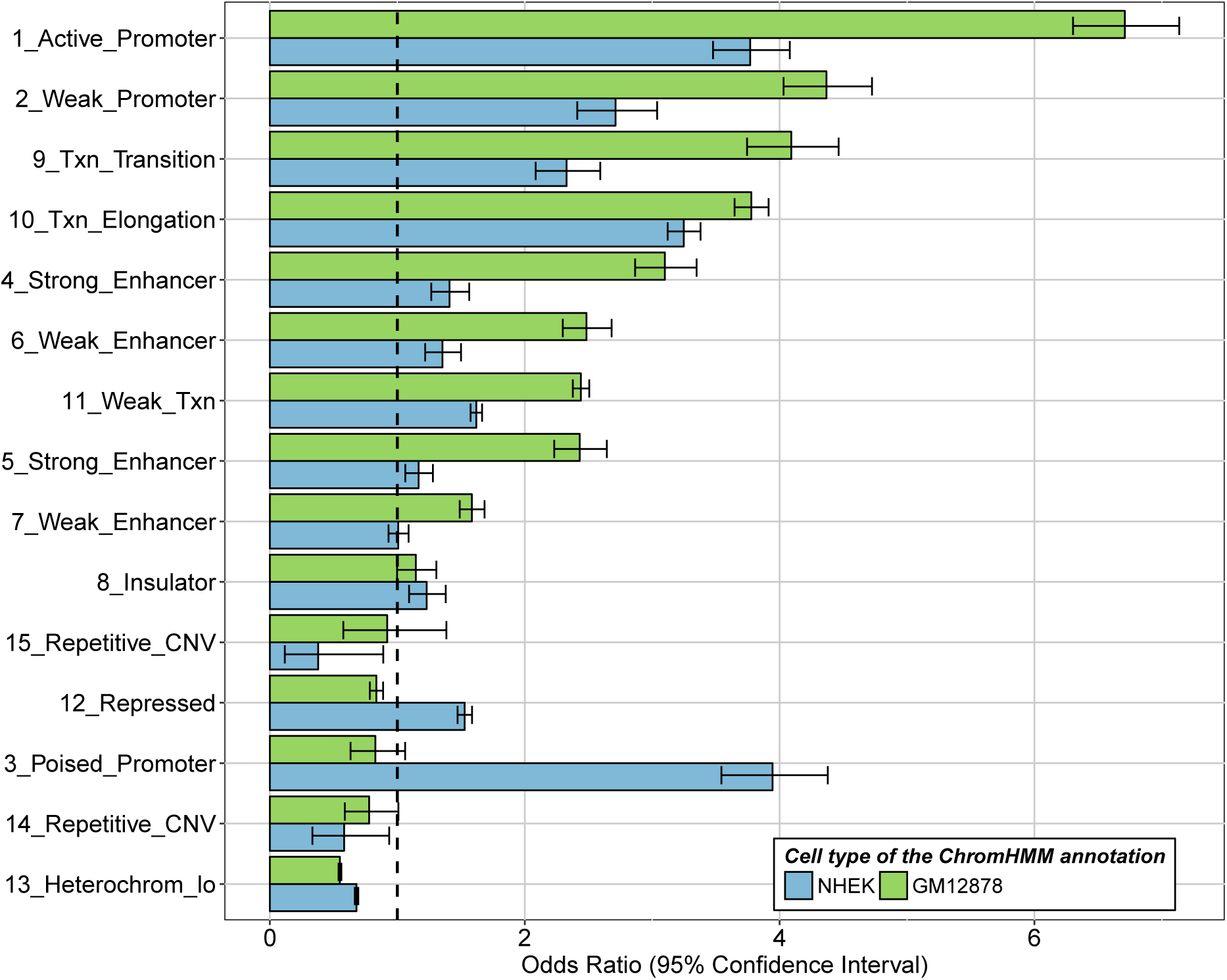
The results of enrichment analysis done performed with all the chromatin states of the relevant cell type were compared with those obtained using the ChromHMM annotation of another cell type (NHEK). For each category, the OR obtained by logistic regression and the corresponding 95% CIs are shown.

**Supplementary figure 6:**
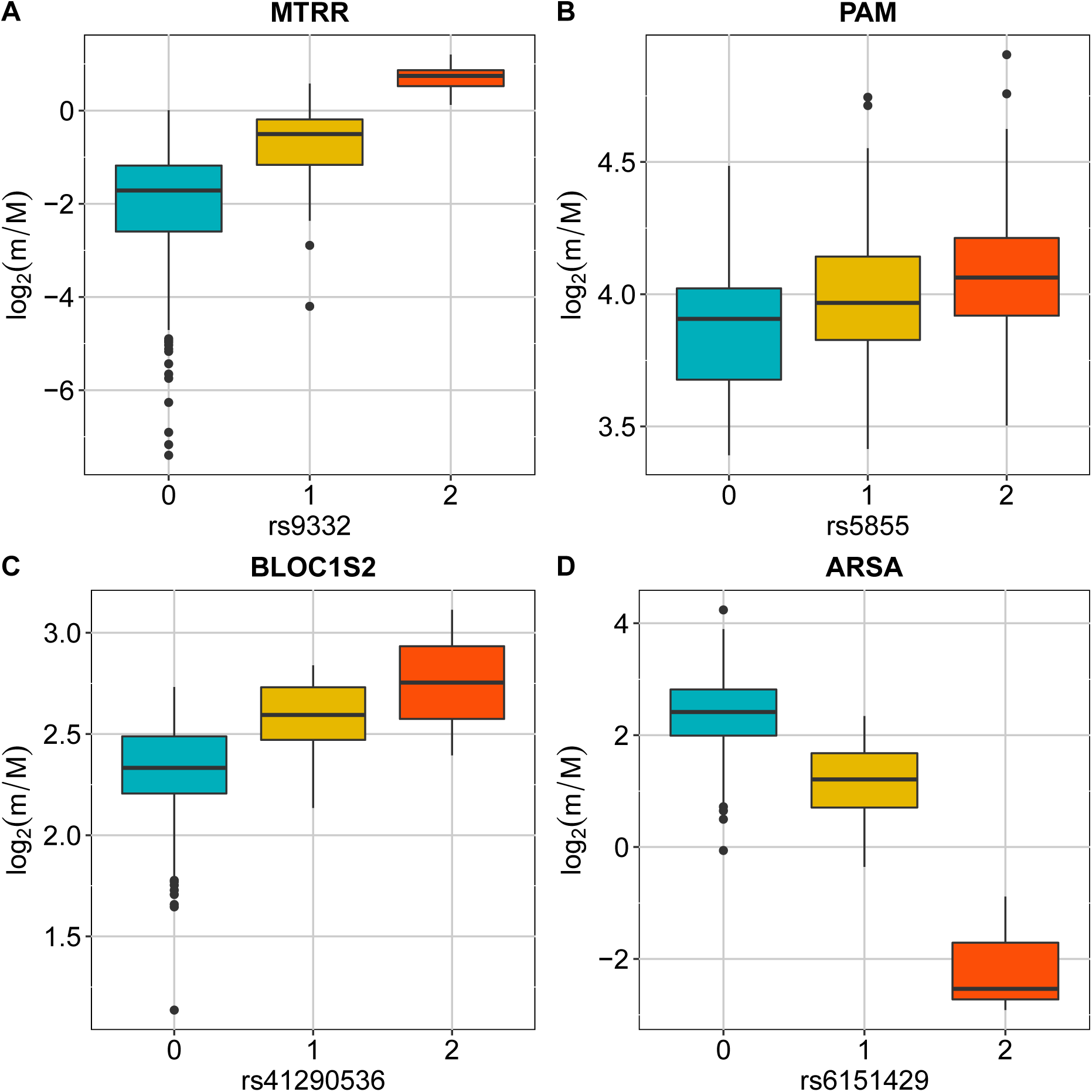
Boxplots showing the variation of the log2-transformed m/M values obtained for MTRR (A), PAM (B), BLOC1S2 (C) and ARSA (D), as a function of the genotype of the individuals for a single genetic variant that falls within the cis-window of the tested gene (rs9332, rs5855, rs41290536 and rs6151429, respectively).

**Supplementary figure 7:**
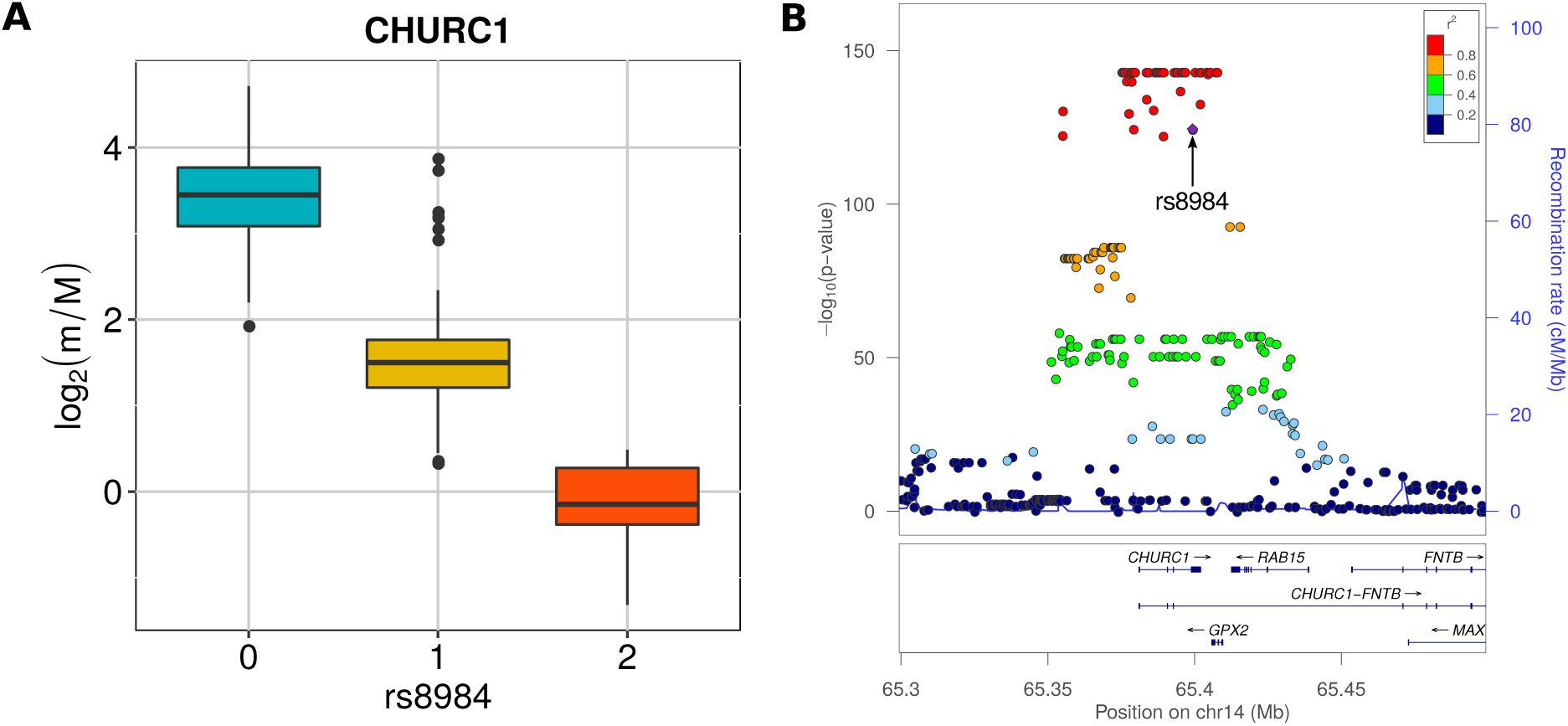
(A) Boxplot showing the variation of log2-transformed m/M values obtained for CHURC1 as a function of the genotype of the individuals for rs8984. (B) LocusZoom plot illustrating the results obtained for CHURC1 in the genomic region around rs8984 (100kb both upstream and downstream its genomic location).

**Supplementary figure 8:**
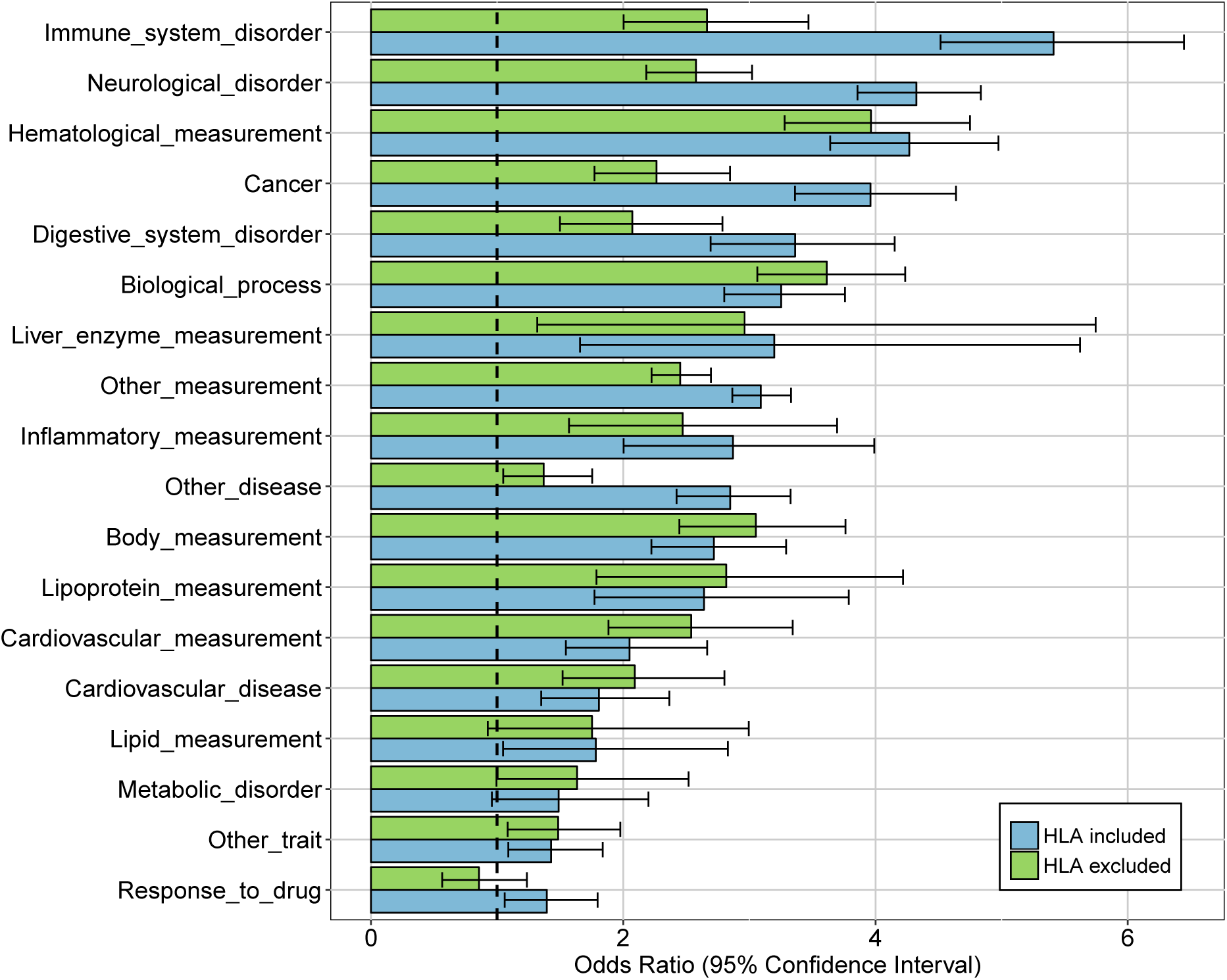
Comparison of the results of the enrichment analyses performed for multiple categories of complex traits considering all the studied genetic variants (HLA included) or after having excluded those that are located within the HLA locus (HLA excluded). For each category, the OR obtained by logistic regression and the corresponding 95% CIs are shown.

**Supplementary figure 9:**
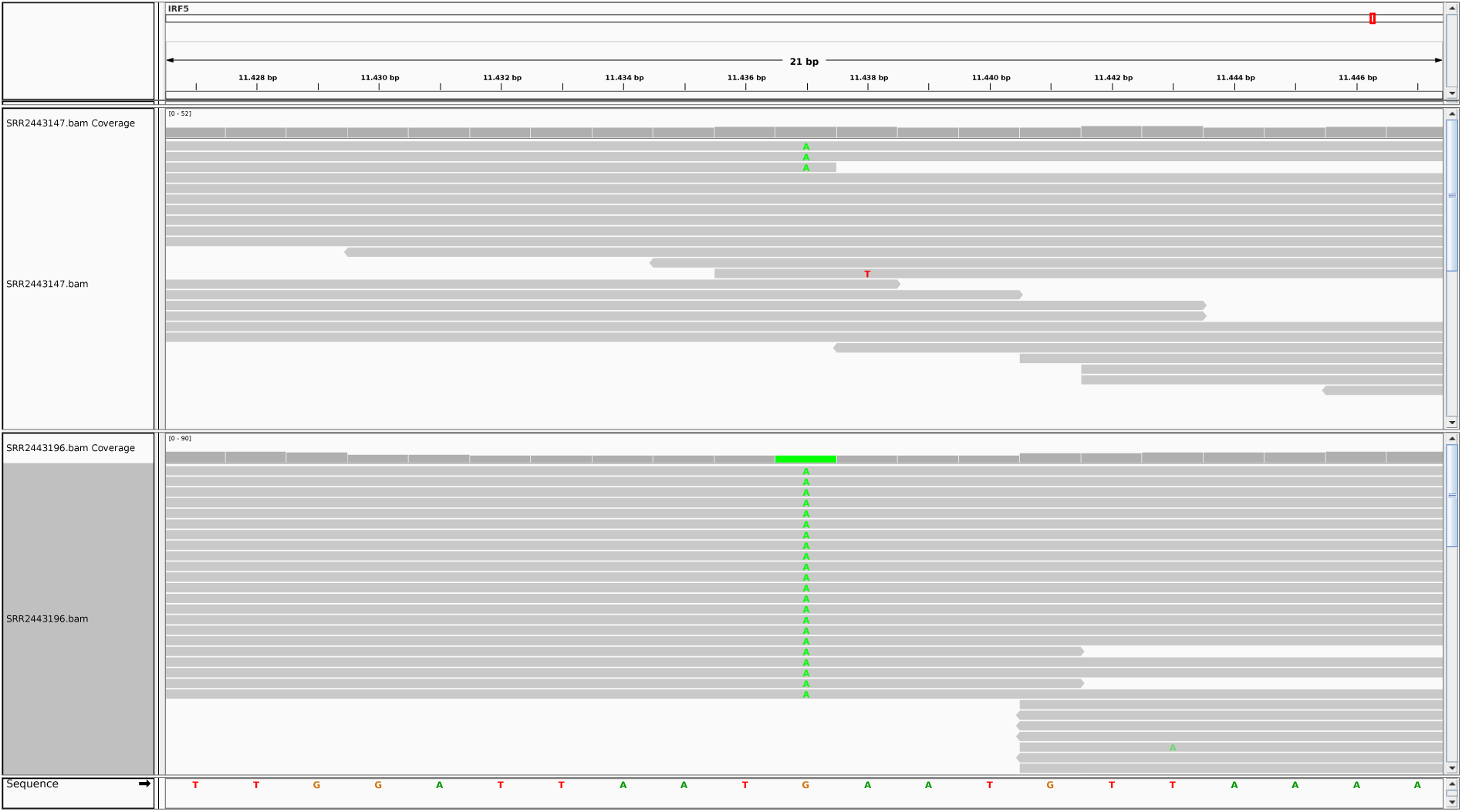
Genotypic information was not available for the SLE patients, therefore their genotype in correspondence of the rs10954213 genetic variant was inferred from RNA-Seq data. In the main analysis, having at least one aligned read with a different nucleotide was considered sufficient to call a heterozygous individual. The figure shows the alignment of RNA-Seq reads in a region around the variant with respect to a reduced genome including only the IRF5 gene and was generated using the Integrative Genomics Viewer (IGV) software [108]. For example, the SRR2443147 sample (top panel) was considered heterozygous, while the SRR2443196 sample (bottom panel) was considered homozygous for the alternative allele.

**Supplementary figure 10:**
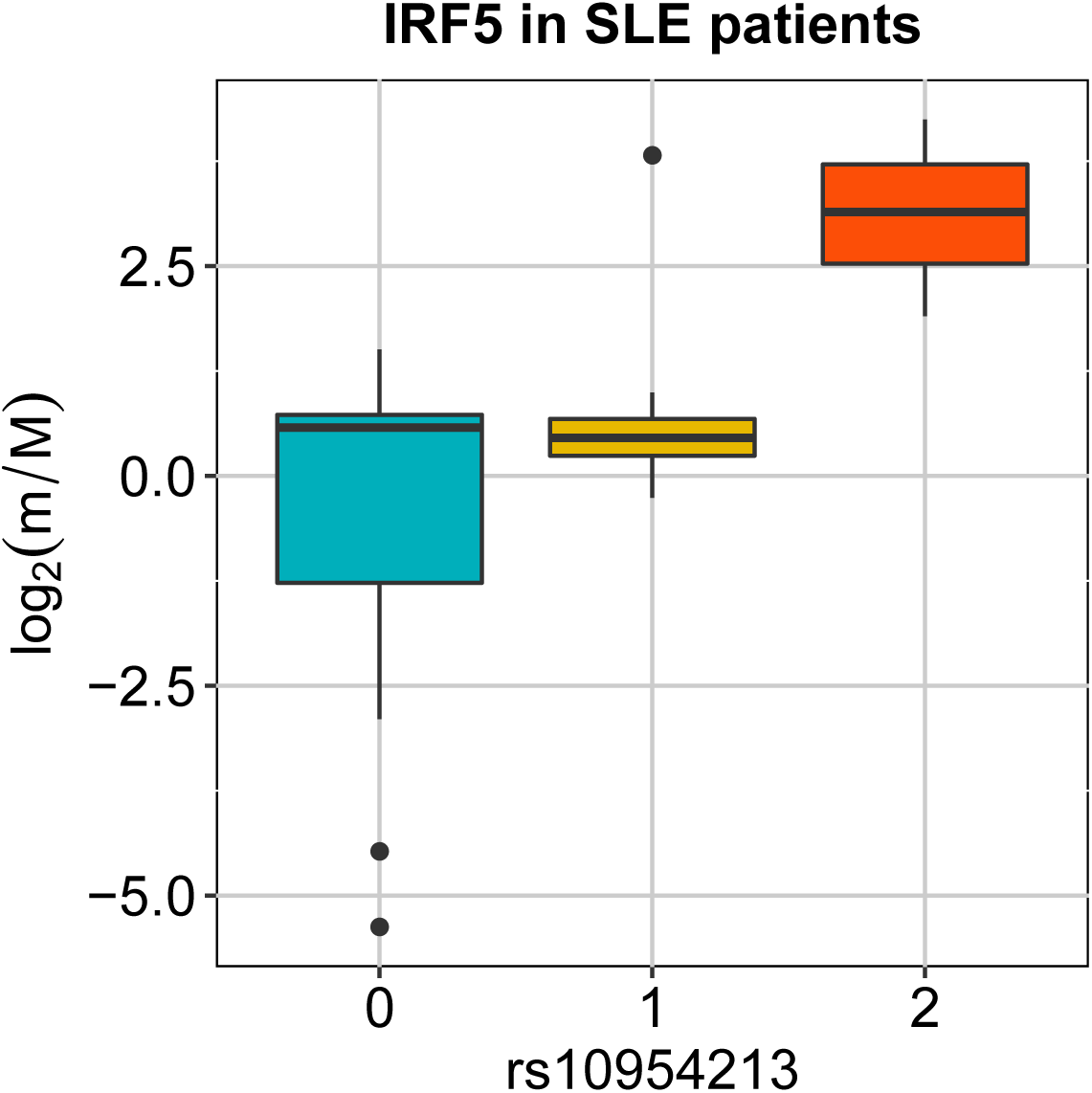
Boxplot showing the variation of the log2-transformed values obtained for IRF5 in SLE patients as a function of the genotype of the individuals when an alternative criterium was adopted to define the genotypic classes.

## Supplementary tables

**Supplementary Table 1:** Enrichment of RBP-altering SNPs among intragenic apaQTL. The table includes the following information for each RNA motif for which a significant enrichment was observed: the identifier of the RNA motif in the CISBP-RNA database (MOTIF ID), the name of the matched RBPs that are expressed in the GM12878 (RBP NAMES), the odds ratio (OR), its 95% confidence interval (95% CI), and the corresponding P-value before and after multiple testing correction by the Benjamini-Hochberg method (P-VALUE and FDR).

| MOTIF ID | RBP NAMES | OR | 95% CI | P-VALUE | FDR |
| --- | --- | --- | --- | --- | --- |
| M016_0.6 | FMR1 | 3.72 | 1.01-11.3 | 0.0278 | 0.266 |
| M025_0.6 | HNRNPC | 1.77 | 1.06-2.83 | 0.0212 | 0.265 |
| M070_0.6 | ENSG00000180771;SRSF2 | 3.83 | 1.55-8.69 | 0.00197 | 0.0521 |
| M075_0.6 | TIA1 | 1.81 | 0.994-3.1 | 0.0388 | 0.327 |
| M081_0.6 | CSDA;YB-1 | 3.9 | 1.03-12.6 | 0.0285 | 0.266 |
| M089_0.6 | HNRNPL | 3.4 | 1.08-9.12 | 0.0217 | 0.265 |
| M122_0.6 | MEX3B;MEX3C;MEX3D | 3.85 | 1.03-12 | 0.0266 | 0.266 |
| M140_0.6 | ENOX1;ENOX2 | 3.16 | 1.36-6.75 | 0.00433 | 0.0778 |
| M145_0.6 | RBM5 | 4.98 | 1.51-14.6 | 0.0044 | 0.0778 |
| M147_0.6 | CNOT4 | 2.11 | 0.947-4.25 | 0.0479 | 0.363 |
| M156_0.6 | TIA1 | 1.81 | 1.05-2.96 | 0.0246 | 0.266 |
| M158_0.6 | HNRNPCL1 | 1.77 | 1.06-2.83 | 0.0212 | 0.265 |
| M160_0.6 | KHDRBS1 | 4.15 | 1.54-10.2 | 0.00271 | 0.0616 |
| M250_0.6 | CSDA | 2.93 | 0.938-7.73 | 0.0411 | 0.327 |
| M256_0.6 | ACO1 | 1.92 | 1.07-3.25 | 0.0209 | 0.265 |
| M291_0.6 | EIF4B | 10.6 | 3.5-33.2 | $2.49 \times 10^{-5}$ | 0.00132 |
| M292_0.6 | EIF4B | 1.98 | 1.32-2.89 | 0.000627 | 0.0249 |
| M320_0.6 | MBNL1;MBNL2;MBNL3 | 1.66 | 1.2-2.25 | 0.00161 | 0.0513 |
| M333_0.6 | SRSF9 | 1.8 | 0.99-3.07 | 0.0397 | 0.327 |
| M344_0.6 | RBMX;RBMXL1;RBMXL2 | 1.77 | 1.4-2.2 | $6.44 \times 10^{-7}$ | $5.12 \times 10^{-5}$ |

**Supplementary Table 2:** Enrichment of GWAS hits for different trait categories among apaQTL. The table includes the following information for each trait category: URI of the trait category in the EFO database (EFO URI) and the associated name (EFO TERM), odds ratio (OR), its 95% confidence interval (95% CI), and the corresponding P-value before and after multiple testing correction by the Benjamini-Hochberg method (P-VALUE and FDR).

| EFO URI | EFO TERM | OR | 95% CI | P-VALUE | FDR |
| --- | --- | --- | --- | --- | --- |
| EFO_0000540 | Immune_system_disorder | 5.41 | 4.52-6.45 | $2.5 \times 10^{-77}$ | $1.5 \times 10^{-76}$ |
| EFO_0000618 | Neurological_disorder | 4.32 | 3.86-4.83 | $2.47 \times 10^{-142}$ | $2.23 \times 10^{-141}$ |
| EFO_0004503 | Hematological_measurement | 4.27 | 3.64-4.98 | $2.89 \times 10^{-74}$ | $1.3 \times 10^{-73}$ |
| EFO_0000616 | Cancer | 3.96 | 3.36-4.64 | $4.15 \times 10^{-63}$ | $1.49 \times 10^{-62}$ |
| EFO_0000405 | Digestive_system_disorder | 3.36 | 2.69-4.15 | $4.34 \times 10^{-28}$ | $9.76 \times 10^{-28}$ |
| GO_0008150 | Biological_process | 3.25 | 2.8-3.76 | $9.69 \times 10^{-56}$ | $2.91 \times 10^{-55}$ |
| EFO_0004582 | Liver_enzyme_measurement | 3.2 | 1.66-5.62 | 0.000167 | 0.000215 |
| EFO_0001444 | Other_measurement | 3.09 | 2.86-3.33 | $1.22 \times 10^{-189}$ | $2.19 \times 10^{-188}$ |
| EFO_0004872 | Inflammatory_measurement | 2.87 | 2-3.99 | $1.77 \times 10^{-09}$ | $3.19 \times 10^{-09}$ |
| EFO_0000408 | Other_disease | 2.85 | 2.42-3.33 | $2.12 \times 10^{-38}$ | $5.46 \times 10^{-38}$ |
| EFO_0004324 | Body_measurement | 2.72 | 2.22-3.29 | $1.59 \times 10^{-23}$ | $3.19 \times 10^{-23}$ |
| EFO_0004732 | Lipoprotein_measurement | 2.64 | 1.77-3.79 | $5.06 \times 10^{-07}$ | $7.58 \times 10^{-07}$ |
| EFO_0004298 | Cardiovascular_measurement | 2.05 | 1.55-2.66 | $2.29 \times 10^{-07}$ | $3.74 \times 10^{-07}$ |
| EFO_0000319 | Cardiovascular_disease | 1.81 | 1.35-2.37 | $3.47 \times 10^{-05}$ | $4.8 \times 10^{-05}$ |
| EFO_0004529 | Lipid_measurement | 1.78 | 1.05-2.83 | 0.0219 | 0.0232 |
| EFO_0000589 | Metabolic_disorder | 1.49 | 0.958-2.2 | 0.0596 | 0.0596 |
| EFO_0000001 | Other_trait | 1.43 | 1.09-1.84 | 0.00751 | 0.00902 |
| GO_0042493 | Response_to_drug | 1.39 | 1.06-1.8 | 0.0132 | 0.0149 |

**Supplementary Table 3:** Enrichment of GWAS hits for different trait categories among apaQTL, after the exclusion of genetic variants within the HLA locus. The table includes the following information for each trait category:URI of the trait category in the EFO database (EFO URI) and the associated name (EFO TERM), odds ratio (OR), its 95% confidence interval (95% CI), and the corresponding P-value before and after multiple testing correction by the Benjamini-Hochberg method (P-VALUE and FDR).

| EFO URI | EFO TERM | OR | 95% CI | P-VALUE | FDR |
| --- | --- | --- | --- | --- | --- |
| EFO_0004503 | Hematological_measurement | 3.96 | 3.28-4.75 | $3.45 \times 10^{-48}$ | $2.07 \times 10^{-47}$ |
| GO_0008150 | Biological_process | 3.61 | 3.06-4.23 | $1.44 \times 10^{-54}$ | $1.29 \times 10^{-53}$ |
| EFO_0004324 | Body_measurement | 3.05 | 2.45-3.76 | $2.99 \times 10^{-24}$ | $1.08 \times 10^{-23}$ |
| EFO_0004582 | Liver_enzyme_measurement | 2.96 | 1.32-5.75 | 0.00338 | 0.00467 |
| EFO_0004732 | Lipoprotein_measurement | 2.82 | 1.79-4.22 | $2.05 \times 10^{-06}$ | $3.99 \times 10^{-06}$ |
| EFO_0000540 | Immune_system_disorder | 2.66 | 2-3.47 | $2.45 \times 10^{-12}$ | $7.36 \times 10^{-12}$ |
| EFO_0000618 | Neurological_disorder | 2.58 | 2.18-3.02 | $3.14 \times 10^{-30}$ | $1.41 \times 10^{-29}$ |
| EFO_0004298 | Cardiovascular_measurement | 2.54 | 1.88-3.34 | $1.83 \times 10^{-10}$ | $4.11 \times 10^{-10}$ |
| EFO_0004872 | Inflammatory_measurement | 2.47 | 1.57-3.7 | $3.14 \times 10^{-05}$ | $4.71 \times 10^{-05}$ |
| EFO_0001444 | Other_measurement | 2.45 | 2.22-2.7 | $5 \times 10^{-75}$ | $9.01 \times 10^{-74}$ |
| EFO_0000616 | Cancer | 2.26 | 1.77-2.85 | $1.3 \times 10^{-11}$ | $3.35 \times 10^{-11}$ |
| EFO_0000319 | Cardiovascular_disease | 2.09 | 1.52-2.8 | $2.22 \times 10^{-06}$ | $3.99 \times 10^{-06}$ |
| EFO_0000405 | Digestive_system_disorder | 2.07 | 1.5-2.79 | $4 \times 10^{-06}$ | $6.55 \times 10^{-06}$ |
| EFO_0004529 | Lipid_measurement | 1.75 | 0.926-3 | 0.0589 | 0.0623 |
| EFO_0000589 | Metabolic_disorder | 1.63 | 0.994-2.52 | 0.0373 | 0.0419 |
| EFO_0000001 | Other_trait | 1.48 | 1.08-1.98 | 0.0099 | 0.0127 |
| EFO_0000408 | Other_disease | 1.37 | 1.05-1.75 | 0.0164 | 0.0197 |
| GO_0042493 | Response_to_drug | 0.856 | 0.565-1.24 | 0.435 | 0.435 |

## Supplementary data

1. **Supplementary File 1: Annotation of gene structures** GTF file with the custom gene annotation that was used for all the analyses.
2. **Supplementary File 2: Annotation of alternative 3′UTR isoforms** GTF file used for the computation of m/M values. For each gene the coordinates of the PRE and POST segments were obtained combining its structure annotation with the poly(A) sites reported by PolyADB_2. In addition the length of each of these segments is reported.
3. **Supplementary File 3: Results of the fitted apaQTL models** For each model we report: the genetic variant identifier according to dpSNP137, as provided into the GEUVADIS dataset (SNP_ID), the NCBI Entrez ID (GENE_ID), the regression coefficient (BETA), the nominal P-value (NOMINAL_PVALUE), the corresponding empirical P-value (EMPIRICAL_PVALUE) and Benjamini-Hochberg corrected empirical P-value (FDR). Multiple files listing the results obtained in each chromosome are publicly available on Mendeley Data [109].
4. **Supplementary File 4: Significant apaQTL models** Table listing the significant models obtained in the apaQTL mapping analysis. For each model we report: the genetic variant identifier (SNP_ID), the NCBI Entrez ID (GENE_ID), the regression coefficient (BETA), the nominal P-value (NOMINAL_PVALUE), the corresponding empirical P-value (EMPIRICAL_PVALUE) and Benjamini-Hochberg corrected empirical P-value (FDR).
5. **Supplementary File 5: Enrichment of trait-specific GWAS hits among apaQTL** The table includes the following information for each GWAS trait for which we found a significant enrichment: the URI of the trait in the EFO database (EFO_URI), the associated name (EFO_TERM) and its parent term in the EFO database (EFO_PARENT_TERM), the odds ratio (OR), its 95% confidence interval (95% CI) and the corresponding P-value before and after multiple testing correction by the Benjamini-Hochberg method (P-VALUE and FDR).
6. **Supplementary File 6: Enrichment of trait-specific GWAS hits among apaQTL, after the exclusion of genetic variants within the HLA locus** The table includes the following information for each GWAS trait for which we found a significant enrichment after the exclusion of genetic variants within the HLA locus: the URI of the trait in the EFO database (EFO_URI), the associated name (EFO_TERM) and its parent term in the EFO database (EFO_PARENT_TERM), the odds ratio (OR), its 95% confidence interval (95% CI) and the corresponding P-value before and after multiple testing correction by the Benjamini-Hochberg method (P-VALUE and FDR).

